# SpatialFusion: A lightweight multimodal foundation model for pathway-informed spatial niche mapping

**DOI:** 10.64898/2026.03.16.712056

**Authors:** Josephine Yates, Mitra Shavakhi, Toni K. Choueiri, Eliezer M. Van Allen, Caroline Uhler

**Author notes:** these authors contributed equally (Eliezer M. Van Allen, Caroline Uhler).

## Abstract

Foundation models enable knowledge transfer across data modalities and tasks, yet foundation models for spatial biology remain in their early stages, largely centered on encoding single-cell representations in spatial context without fully integrating transcriptomic and morphological information to delineate functional niches. Here we introduce SpatialFusion, a lightweight multimodal foundation model that identifies biologically coherent microenvironments defined by distinct pathway activation patterns rather than spatial proximity alone. SpatialFusion integrates paired histopathology, gene expression, and inferred pathway activity into a unified representation. Compared with two specialist niche-detection methods and four spatial foundation models, SpatialFusion performs competitively and consistently resolves fine-grained spatial niches with unique pathway-level signatures. Applying the model to two Visium HD cohorts uncovered a pre-malignant niche in morphologically normal mucosa adjacent to colorectal tumors and revealed distinct malignant microenvironments in non-small cell lung cancer that were predictive of tumor stage. Overall, SpatialFusion offers a versatile framework for multimodal spatial analysis, enabling the discovery of new morpho-molecular niches with significant biological and clinical relevance.

## Introduction

Foundation models have recently transformed computational biology in domains like protein structure prediction and single-cell biology by enabling the transfer of knowledge across diverse data modalities and tasks. Trained on large and heterogeneous datasets, these deep learning architectures capture high-dimensional biological structure, often exhibiting emergent capabilities such as batch correction, without explicit supervision. In single-cell genomics, models such as scGPT^1^ and Geneformer^2^ have demonstrated powerful generalization across cell types and tissues, while in digital pathology, foundation models including UNI^3^ and H-optimus (https://huggingface.co/bioptimus/H-optimus-1) have shown similar versatility in the analysis of hematoxylin and eosin (H&E)–stained images.

The advent of spatially resolved transcriptomic technologies has introduced a new dimension to molecular profiling by coupling gene expression measurements with the spatial architecture of tissues. These datasets, often at single-cell or near–single-cell resolution, enable detailed analysis of cellular context, microenvironmental interactions, and tissue organization. Spatial information is particularly critical for understanding disease progression, therapeutic resistance, and tumor–immune dynamics in cancer^4,5^. In this context, spatial niches have been closely associated with disease features and progression^6–8^. Capturing these functional ecosystems requires models that integrate spatial and molecular information to distinguish regions based on pathway activation patterns rather than cell identity alone. This level of mechanistic insight is especially important in cancer, where malignant cells rewire signaling pathways and remodel their microenvironment through immune and stromal interactions; identifying niches defined jointly by spatial organization and pathway activity can therefore reveal the ecological structures that support tumor growth, immune evasion, and therapeutic response and resistance.

Specialized computational methods such as NicheCompass^9^ and BANKSY^10^ have been developed to identify spatial niches in high-gene-coverage spatial transcriptomics (ST) data (e.g., Visium and Xenium). Although effective for niche discovery, these models are not pretrained on large, diverse corpora of tissues and therefore do not benefit from the broad biological coverage, batch-robustness, or emergent representations characteristic of foundation models. Moreover, these approaches cannot operate on H&E images alone, limiting their applicability to settings without paired transcriptomic data. In parallel, spatial foundation models are beginning to emerge but remain in an early stage. Recent approaches such as NicheFormer^11^, scGPT-spatial^12^, HEIST^13^, Novae^14^, and OmiCLIP^15^, have begun extending transformer and contrastive architectures to spatial omics, yet each has notable constraints. NicheFormer and scGPT-spatial produce transformer-based single-cell embeddings: NicheFormer is trained on spatial datasets but does not incorporate spatial coordinates into the architecture, whereas scGPT-spatial predicts a cell’s gene expression using its neighbors and therefore encodes limited spatial context, but outputs single-cell rather than neighborhood-level representations. HEIST uses graph attention networks to integrate gene regulatory and spatial information, but operates solely on transcriptomics and remains unimodal. Novae is a graph-based foundation model for spatial transcriptomics that learns self-supervised cell representations within their spatial context and performs zero-shot domain inference across tissues and technologies. OmiCLIP jointly embeds histopathology and transcriptomic data via contrastive learning on Visium, but functions at the spot level and, like NicheFormer, does not explicitly encode spatial coordinates during training. As a result, most current spatial foundation models incorporate minimal explicit spatial reasoning, prioritize cell-level embeddings, and do not fully integrate transcriptomic and morphological information into coherent, biologically interpretable niches. Moreover, none explicitly identify multicellular niches defined by coordinated pathway activation across cell types, a key requirement in contexts such as cancer, where dysregulated signaling and microenvironmental crosstalk shape disease progression.

To address these limitations, we developed SpatialFusion, a lightweight multimodal foundation model that integrates paired histopathological and transcriptomic data with pathway activity scores to produce biologically grounded spatial neighborhood embeddings at single-cell resolution. Its lightweight design stems from the use of precomputed foundation model embeddings, yielding low memory and computational requirements and eliminating the need for specialized acceleration hardware. SpatialFusion combines the advantages of foundation models (pretraining on a large, diverse multi-tissue corpus) with the niche-centric design of specialist methods. SpatialFusion outperforms existing models in niche rediscovery across multiple tissue types and performs competitively in zero-shot evaluations. We further demonstrate the versatility of SpatialFusion by applying it to Visium HD cohorts, where it uncovers pre-malignant niches in colorectal cancer and identifies spatial niches associated with lymph node metastasis in non-small cell lung cancer.

## Results

### SpatialFusion integrates histopathology and transcriptomics to learn biologically-meaningful multimodal representations of spatial niches

SpatialFusion is a deep learning framework that integrates spatial transcriptomics (ST) data with hematoxylin and eosin (H&E) histopathology to generate joint embeddings of cellular neighborhoods, which are subsequently grouped into spatial niches. Operating at single-cell resolution, the method can be applied to paired H&E and ST datasets or to whole-slide H&E images alone. By combining molecular and morphological information, SpatialFusion captures coordinated patterns of tissue organization and gene expression.

We define a spatial niche as a reproducible, functionally specialized microenvironment characterized by coordinated transcriptional programs and structured cell–cell interactions, in contrast to a neighborhood, which reflects only spatial proximity without implying functional organization. Spatial niches are therefore distinguished by molecular coordination across cell types rather than by cellular composition alone. This biological prior is embedded directly into the model design: SpatialFusion is explicitly trained to encode pathway-level activation signatures in its latent space, enabling the identification of microenvironments defined by distinct functional states. As a result, the method recovers biologically meaningful niches that integrate morphology, signaling activity, and microenvironmental context.

SpatialFusion is trained on a subset of Xenium ST experiments from the HEST1-k dataset^16^ (51 samples, approximately 11 million cells), spanning 12 tissue types across healthy, diseased, and cancerous states (Supplementary Fig.1). The architecture consists of three main modules: a unimodal encoding module, a multimodal alignment module, and a spatial encoding module (Fig. 1a).

**Figure 1:**
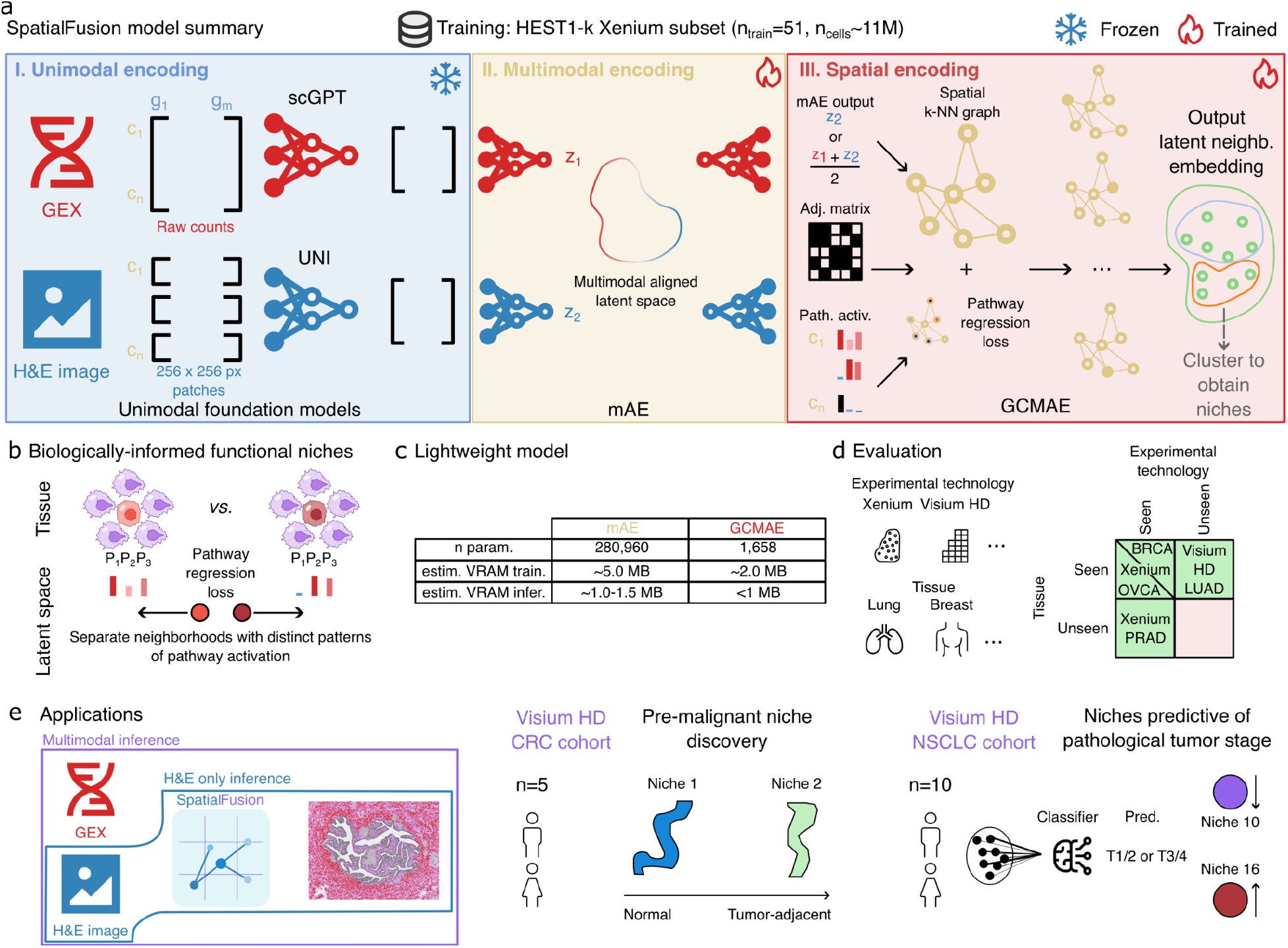
Overview of the SpatialFusion framework. **a**, Schematic of the SpatialFusion architecture. Paired gene expression counts (GEX) and hematoxylin and eosin (H&E) image patches are encoded using frozen foundation models, scGPT and UNI, to generate low-dimensional embeddings (z). These embeddings are aligned through a trained multimodal autoencoder (mAE) and then used as node features in a trained graph-convolutional masked autoencoder (GCMAE). Tissues are represented as spatial graphs using the spatial adjacency matrix, in which nodes correspond to cells and edges connect the 30 nearest neighbors. An auxiliary pathway regression loss, used to predict the pathway activation pattern provided as input to the model, is added to the standard GCMAE objectives to encourage the latent space to capture pathway activation patterns in the latent neighborhood embedding, which is the output of the model. Niches can be obtained by clustering these neighborhood embeddings. **b**, The pathway regression loss enables SpatialFusion to distinguish spatial neighborhoods with similar cell-type composition but divergent pathway activity. Two cells with similar neighborhoods in terms of cell type composition but different pathway activation patterns (pathways P1, P2, and P3) will be separated in the latent space. **c**, SpatialFusion is lightweight, containing fewer than 300,000 learnable parameters and requiring under 5 MB of VRAM for training and under 2 MB for inference. **d**, The model is trained exclusively on Xenium spatial transcriptomics data yet generalizes across tissue types and experimental platforms, including unseen technologies such as Visium HD. BRCA: breast cancer, OVCA: ovarian cancer, PRAD: prostate cancer, LUAD: lung adenocarcinoma. **e**, SpatialFusion can be applied to paired H&E and spatial transcriptomics datasets for multimodal inference or to H&E whole-slide images for image-only inference. Applications across cancer cohorts demonstrate its ability to discover biologically meaningful niches, including a pre-malignant niche within morphologically normal mucosa in colorectal cancer (CRC) and niches predictive of T stage in non-small cell lung cancer (NSCLC). Abbreviations: GEX: gene expression; H&E: hematoxylin and eosin; px: pixel; adj. matrix: adjacency matrix; path. activ.: pathway activation; VRAM: video random access memory; OVCA: ovarian cancer; BRCA: breast cancer; PRAD: prostate cancer; LUAD: lung adenocarcinoma; CRC: colorectal cancer; NSCLC: non-small cell lung cancer; T1/2/3/4: pathological tumor stage 1/2/3/4.

In the unimodal encoding module, each cell is represented by raw gene expression counts and a 256 × 256-pixel H&E patch. These inputs are embedded using frozen foundation models, scGPT^1^ for the transcriptomic input and UNI2^3^ for the imaging input, chosen for their strong performance in recent benchmarks^17,18^. This initialization yields technology-agnostic, batch-corrected representations for each modality.

The multimodal alignment module uses a multimodal autoencoder (mAE) to align the paired transcriptomic and histological embeddings in a shared latent space, producing low-dimensional representations at single-cell resolution that capture concordant biological variation. Depending on the application, SpatialFusion can use the joint multimodal embedding (an average of the transcriptomic and imaging representations) or the H&E-only embedding for image-only inference.

The spatial encoding module models the tissue as a k-nearest-neighbor graph in which nodes correspond to cells and edges capture spatial proximity. A graph convolutional masked autoencoder (GCMAE) is trained on this graph to learn a latent embedding for each cell’s local neighborhood. To encourage the model to capture functional microenvironments rather than spatial adjacency alone, we introduce an auxiliary pathway regression loss. Pathway activity scores are estimated using PROGENy^19^ for ten pathways relevant to cancer biology, including EGFR, androgen, estrogen, JAK–STAT, VEGF, MAPK, PI3K, TGFβ, NFκB, and TGFA. These scores are incorporated as auxiliary targets to enforce pathway-level structure in the latent space, enabling the model to differentiate spatial neighborhoods with similar cell type composition but distinct functional states (Fig. 1b).

The model outputs a latent embedding for each cell that summarizes the local neighborhood centered on that cell. Clustering these by embeddings yields groups of recurrent spatial contexts, which we define as niches (Fig. 1a).

SpatialFusion is designed to be computationally lightweight, with fewer than 300,000 learnable parameters (~280,000 in the mAE and ~2,000 in the GCMAE) and memory requirements under 5 MB VRAM for training and under 2 MB for inference (Fig. 1c, Supplementary Methods). The only computationally demanding step is the one-time extraction of scGPT and UNI embeddings during preprocessing. Once these embeddings are precomputed, SpatialFusion can be executed on a standard GPU-equipped laptop.

We evaluate SpatialFusion across multiple settings that represent major challenges in the field and key use cases for discovery: in tissue types included during training, in entirely unseen tissue types, and on an unseen experimental platform (Visium HD) (Fig. 1d). Because SpatialFusion is multimodal, it supports both paired H&E and ST inference and H&E-only inference, enabling application to large cohorts lacking spatial transcriptomics (Fig. 1e). To showcase SpatialFusion’s usefulness, we apply it to two Visium HD cancer cohorts: a public colorectal cancer (CRC) dataset and an in-house non-small cell lung cancer (NSCLC) dataset (Fig. 1e).

A key feature of SpatialFusion is its reliance on frozen foundation models as unimodal encoders, which offers two major advantages: (i) training remains lightweight because the computational bottleneck is limited to a single round of foundation-model inference; and (ii) despite the limited availability of paired H&E and single-cell–resolution spatial data for training, SpatialFusion inherits biologically meaningful, batch-robust representations learned from millions of cells and image patches in the foundation models. To evaluate this design, we compared the multimodal autoencoder with a baseline model using an MLP encoder for gene expression and a fine-tuned ImageNet-pretrained ResNet for images (Supplementary Fig.1). Because this stage involves no spatial information, we assessed embedding quality using single-cell metrics. Biological signal preservation was evaluated using a logistic regression classifier and balanced accuracy (BAC), while batch correction was quantified using the batch average silhouette width (bASW)^20^. SpatialFusion outperformed the baseline by 16 percent in BAC (0.55 to 0.71) and by 9 percent in bASW (0.86 to 0.95), indicating improved biological fidelity and reduced batch effects (Supplementary Fig.1).

A second innovation is the pathway regression loss added in the GCMAE. Ridge regression analysis showed that this objective improves organization of the latent space with respect to pathway activation, increasing average R^2^ across PROGENy pathways by 5 percent (range: 2.4 to 9.4 percent) (Supplementary Table 1). PCA confirmed clearer pathway separation in models trained with this loss (Supplementary. Fig. 1).

In summary, SpatialFusion fuses molecular and morphological information to generate biologically grounded representations of spatial niches. Through pretrained encoders, multimodal alignment, and pathway-informed spatial learning, the model jointly captures cellular composition, morphological context, and functional signaling. By design, SpatialFusion incorporates key strengths of both spatial foundation models and specialist niche-discovery approaches while addressing their limitations: it leverages large-scale pretraining, learns neighborhood-level rather than single-cell embeddings, integrates H&E and transcriptomic modalities, and explicitly encodes pathway activation patterns. This architecture enables the discovery of distinct, functionally meaningful niches, particularly in cancer.

### SpatialFusion outperforms existing methods in rediscovery of niches in ovarian cancer tissue

The latent embeddings produced by SpatialFusion can be clustered to identify groups of cells that share similar microenvironmental, morphological, and pathway activation characteristics, which we define as spatial niches. To benchmark model performance, we evaluate its capability to rediscover pathologist-annotated niches that served as ground truth for spatial niche identities. Thus, the model was first evaluated on an unseen Xenium ovarian cancer (OVCA) sample comprising approximately 5,000 genes with pathologist-provided spatial annotations and predefined cell-type labels (Fig. 2a–b, Supplementary Fig.2).

**Figure 2:**
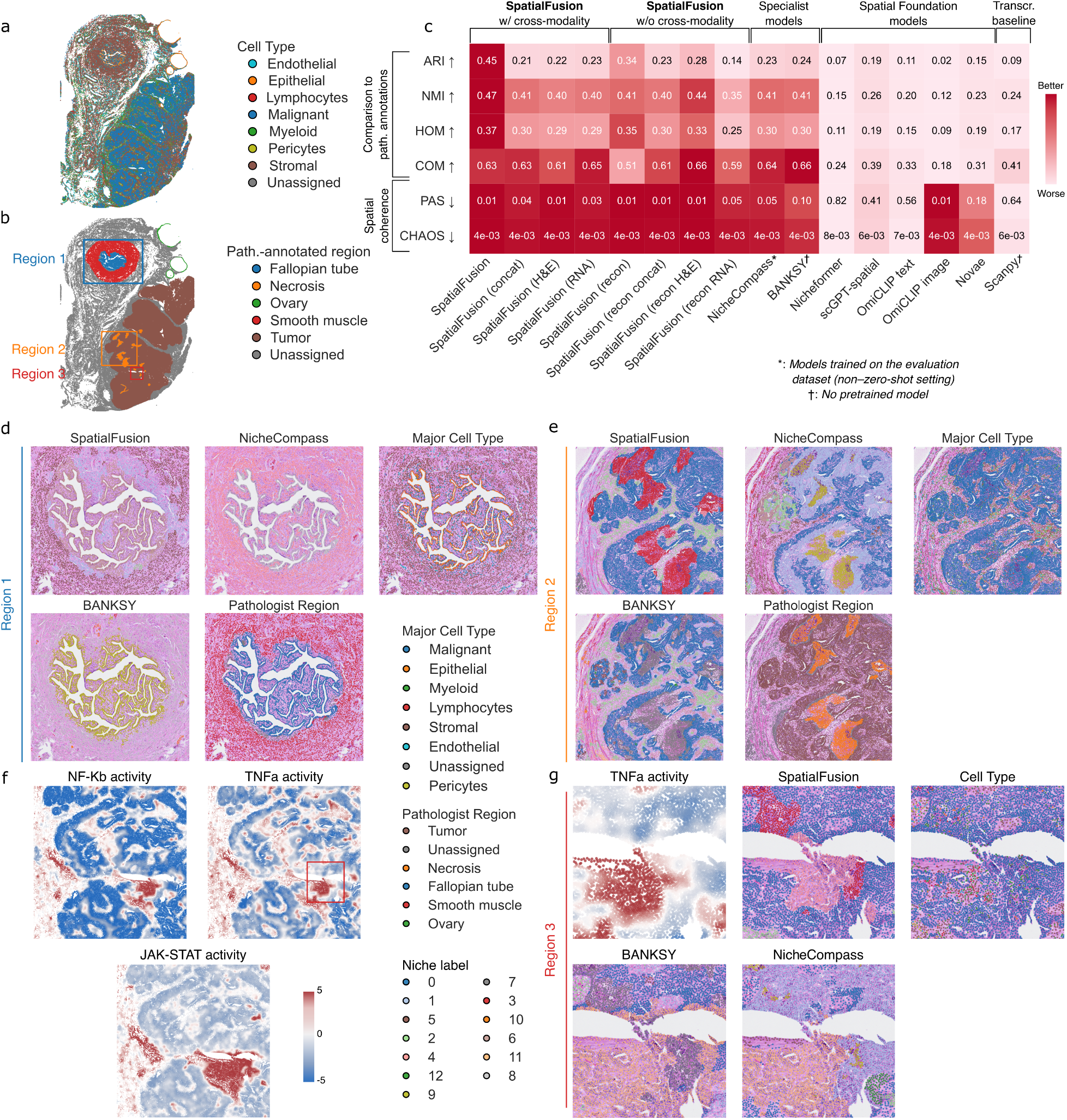
SpatialFusion benchmarked on an ovarian cancer (OVCA) Xenium 5k sample. **a**, Expert-annotated cell types displayed in spatial coordinates on the OVCA sample section. **b**, Pathologist-defined regions, with three regions of interest highlighted. **c**, Benchmark comparing SpatialFusion with specialist niche-detection methods, spatial foundation models, and a transcriptomic baseline. All models were applied to the OVCA sample with default parameters, and resulting embeddings were clustered using Leiden clustering (n = 13). We report the SDMBench^21^ metrics: Adjusted Rand Index (ARI), Normalized Mutual Information (NMI), homogeneity (HOM), and completeness (COM), which quantify agreement with pathologist annotations; and Percentage of Abnormal Spots (PAS) and spatial chaos (CHAOS), which assess spatial coherence. NicheCompass does not provide a pretrained model and was therefore trained directly on the OVCA evaluation dataset; reported metrics thus reflect performance on data used for training. BANKSY and Scanpy do not require training. We also evaluate alternative SpatialFusion configurations, including a model without cross-modality reconstruction (SpatialFusion without cross-modality), averaged or concatenated embeddings, and unimodal RNA-only or H&E-only variants. **d**, Zoomed-in view of region 1 in (b) containing a fallopian tube encircled by smooth muscle. Shown are SpatialFusion, NicheCompass, and BANKSY niches alongside cell-type and pathologist annotations overlaid on the H&E image. **e**, Zoom of region 2 containing tumor and necrotic tissue, with niches from SpatialFusion, NicheCompass, and BANKSY compared to cell-type and pathologist annotations. **f**, Spatial distribution of PROGENy-estimated NF-κB, TNFα, and JAK–STAT pathway activation, highlighting a region with elevated immune signaling (region 3). **g**, Zoom of region 3 showing high NF-κB, TNFα, and JAK–STAT activity. Shown are SpatialFusion, NicheCompass, and BANKSY niches, as well as cell types and TNFα activation overlaid on the H&E image.

We compared SpatialFusion with established methods designed for this task, including NicheCompass^9^ and BANKSY^10^ (Fig. 2c). BANKSY embeds each cell into a combined “product space” that integrates its own molecular profile with the average profile of its spatial neighbors, followed by clustering that augmented space. NicheCompass constructs a spatial graph of cellular neighborhoods and uses a graph variational autoencoder to learn embeddings that capture intercellular communication programs.

We also compared SpatialFusion with spatial foundation models introduced earlier, including NicheFormer^11^, scGPT-spatial^12^, Novae^14^, and OmiCLIP^15^ (Fig. 2c). Because OmiCLIP can encode both transcriptomic and histological data, we evaluated its performance separately for the transcriptomic-based and image-based embeddings. As a baseline, we included unsupervised Leiden clustering performed directly in transcriptomic space using Scanpy, without incorporating spatial information. All external models were applied with their default settings, as showcased in tutorials (Methods). For all models, we performed Leiden clustering (n = 13).

We assessed performance using four clustering metrics (adjusted Rand index (ARI), normalized mutual information (NMI), homogeneity (HOM), and completeness (COM)) and two spatial coherence metrics (percentage of abnormal spots (PAS) and spatial chaos (CHAOS)) (Methods).

In this benchmark, SpatialFusion outperformed all other methods across all evaluation metrics except completeness (COM), where the difference was small (Δ = 0.01–0.02) (Fig. 2c, Supplementary Fig. 2). The slight reduction in completeness likely reflects over-segmentation of certain ground-truth classes, producing purer but more fragmented clusters. Importantly, SpatialFusion and the other foundation models were evaluated in a zero-shot setting, where pretrained models were directly applied to the ovarian cancer (OVCA) dataset without retraining. In contrast, NicheCompass, which lacks a pretrained version, was trained on the OVCA data, while BANKSY and Scanpy were applied directly without training. Among the evaluated models, NicheCompass and BANKSY achieved the second-best overall performance. Some methods, such as OmiCLIP (image), failed to produce biologically meaningful embeddings, instead clustering spatially contiguous patches (Supplementary Fig. 2). Similarly, other spatial foundation models primarily grouped cells by cell type rather than by spatial niche, consistent with their design focus on encoding single-cell representations in a spatial context rather than modeling multicellular ecosystems; this was evident from their higher ARI with cell-type annotations relative to niche assignments (Supplementary Fig. 2). This explains their comparable performance to the transcriptomic-only baseline in this benchmark. To confirm the robustness of these findings, we repeated the benchmark using different numbers of clusters, and the relative ranking and performance trends among methods remained stable (Supplementary Fig. 2).

We next examined alternative configurations of SpatialFusion, including a model trained without cross-modality reconstruction (SpatialFusion w/o cross-modality), models that combined embeddings by averaging or concatenation, and unimodal variants that used only the transcriptomic or H&E branch. Across all metrics except COM, the full SpatialFusion model achieved the highest performance, confirming the advantage of jointly integrating both modalities through cross-modal alignment (Fig. 2c).

To better understand model behavior, we compared the clusters identified by SpatialFusion (multimodal and H&E variants) with those from NicheCompass and BANKSY, the next best-performing methods, against pathologist annotations using confusion matrices (Supplementary Fig. 2). SpatialFusion more accurately rediscovered annotated regions, particularly the fallopian tube, necrotic zones, and tumor areas. In the fallopian tube region (Region 1), NicheCompass predominantly captured major cell type boundaries (epithelial versus non-epithelial), whereas SpatialFusion delineated the entire pathologist-defined region, explaining its superior performance (Fig. 2d). In the tumor region (Region 2), SpatialFusion captured necrotic regions more precisely and represented the tumor mass as a coherent structure, unlike NicheCompass and BANKSY, which produced more fragmented clusters (Fig. 2e).

We also leveraged the pathway activation pattern inference capacity of SpatialFusion to derive biologically interpretable features from this tumor. Within the tumor core (Region 3), we observed a region with high activation of immune-related pathways, including NF-κB, TNFα, and JAK–STAT (Fig. 2f, Supplementary Fig. 2). Closer inspection revealed a lymphoid aggregate, with lymphoid and myeloid cells forming a ring around malignant cells (Fig. 2g). SpatialFusion precisely captured this immune-enriched niche, whereas NicheCompass and BANKSY showed less spatial specificity in this area. These findings highlight SpatialFusion’s ability to resolve fine-grained spatial niches characterized by coordinated pathway activity, enabled by the pathway regression loss used in the model, features that may be overlooked by other approaches.

### SpatialFusion rediscovers meaningful niches in breast cancer

To further assess SpatialFusion’s robustness across tissue types and disease contexts, we evaluated its performance on an additional tissue, breast cancer, and compared its variants with specialist niche-discovery methods, spatial foundation models, and a transcriptomic clustering baseline for rediscovering pathologist-annotated regions. In addition to the multimodal and H&E-only inference modes, we included a fine-tuned version of SpatialFusion, in which the multimodal autoencoder (mAE) and GCMAE were pretrained on the same dataset used for evaluation but without access to the pathologist annotations. This configuration adapts the embeddings to the specific tissue while maintaining unsupervised training, thereby providing a fairer comparison to NicheCompass, which is trained on the evaluation data.

We tested all models on a spatially annotated breast cancer (BRCA) Xenium sample (n=300 genes) containing invasive carcinoma and ductal carcinoma in situ (DCIS) regions (Fig. 3a–b, Supplementary Fig. 3). The annotations include invasive lesions and two distinct DCIS subregions (DCIS1 and DCIS2). In this setting, the zero-shot SpatialFusion variants ranked second to NicheCompass in NMI, HOM, and COM, but achieved the highest spatial coherence scores (Fig. 3c, Supplementary Fig. 3). Performance was comparable to BANKSY, with SpatialFusion outperforming it in ARI, HOM, and spatial metrics, but underperforming in NMI and COM. The fine-tuned version of SpatialFusion improved across all metrics relative to the zero-shot model, reaching performance similar to NicheCompass, though still weaker in NMI, HOM, and COM. The Scanpy baseline also performed competitively, likely reflecting the strong correspondence between cell-type composition and pathological boundaries in this sample.

**Figure 3:**
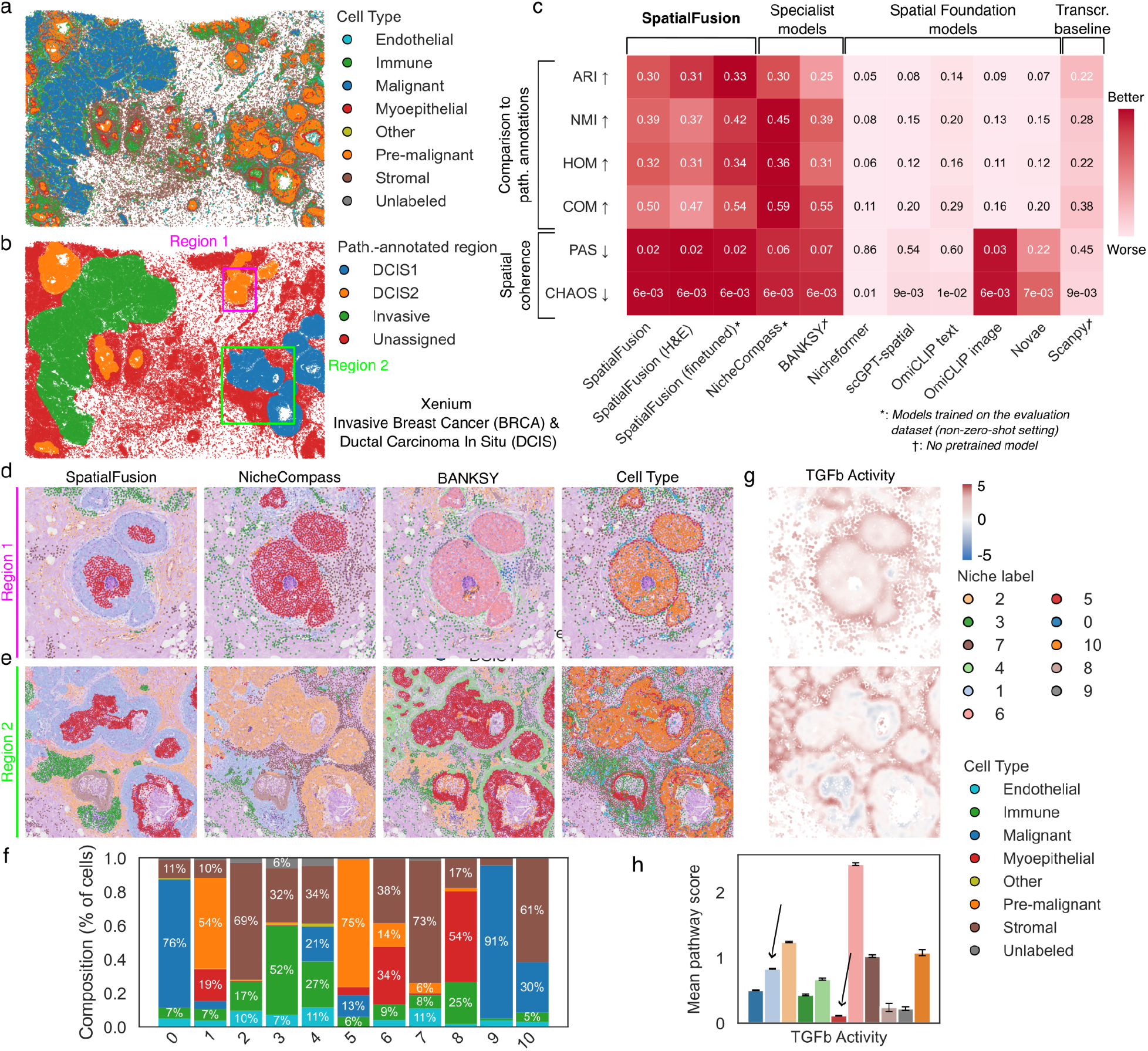
SpatialFusion identifies biologically meaningful niches in invasive breast cancer (BRCA) and ductal carcinoma in situ (DCIS). **a**, Expert-annotated cell types displayed in spatial coordinates on the BRCA sample section. **b**, Pathologist-defined regions, with two regions of interest highlighted. **c**, Benchmark comparing SpatialFusion with specialist niche-detection methods, spatial foundation models, and a transcriptomic baseline, analogous to Fig. 2c. All models were applied to the BRCA sample with default parameters, and embeddings were clustered using Leiden clustering (n = 11). In addition to the zero-shot model, a fine-tuned SpatialFusion variant pretrained on this sample (without using pathologist labels) was evaluated. **d–e**, Zoomed views of **d**, DCIS2 and **e**, DCIS1 regions. Shown are niches identified by SpatialFusion, NicheCompass, and BANKSY, alongside cell-type annotations overlaid on the H&E image. **f**, Cell-type composition of SpatialFusion-derived niches. **g**, PROGENy-estimated TGFβ pathway activation over the same regions shown in panels **d** and **e. h**, Mean TGFβ activation scores for each SpatialFusion-defined niche.

Confusion matrix comparisons revealed that SpatialFusion subdivided the DCIS1 and DCIS2 regions into distinct edge and core compartments (Supplementary Fig. 3). The edge niche (niche 1) corresponded to a boundary enriched in DCIS and myoepithelial cells, while the core niche (niche 5) consisted predominantly of pre-malignant cells (Fig. 3d–f). This boundary was captured by SpatialFusion and BANKSY but not by NicheCompass. The myoepithelial barrier surrounding DCIS lesions is known to regulate the transition to invasive carcinoma and represents a key structural feature of breast cancer pathophysiology^22,23^. Pathway analysis showed that niche 5 had lower TGF-β activity than niche 1 (Fig. 3g–h), in line with reports that TGF-β signaling in the myoepithelial compartment promotes the invasive progression of DCIS^24–26^ Discrepancies between SpatialFusion and pathologist annotations therefore appear biologically meaningful rather than artifactual. These findings further underscore SpatialFusion’s ability to resolve spatial niches that capture both tissue morphology and pathway-level signaling dynamics.

### SpatialFusion recovers niches in unseen tissue and unseen technology

We next evaluated the generalization ability of SpatialFusion in challenging settings involving completely unseen tissues and technologies, that is, tissue types and experimental platforms not encountered during training. Specifically, the model has not been exposed to any prostate tissue or Visium HD data during training; hence, we applied the model to a pathologist-annotated (M.S.) prostate cancer sample and to a lung adenocarcinoma (LUAD) Visium HD dataset processed to single-cell resolution using bin2cell^22^ (Methods);

In the prostate cancer sample (Xenium, number of genes ~10k), all models struggled, likely due to high cellular heterogeneity, with ARI values ranging from 0 to 0.21 and homogeneity from 0.01 to 0.16 (Fig. 4a–c). Some models, such as NicheFormer and BANKSY, failed to produce meaningful clusters under default parameters (Supplementary Fig. 4). Interestingly, OmiCLIP (image) performed best in this setting, while the SpatialFusion (H&E) variant ranked second, narrowly surpassing NicheCompass. The multimodal SpatialFusion variant performed slightly below NicheCompass, but fine-tuning substantially improved performance, yielding results comparable to OmiCLIP (higher COM, lower ARI, and similar values across other metrics). These results indicate that finetuning markedly enhances SpatialFusion’s performance on unseen tissue, although even the zero-shot H&E variant remains competitive despite never having encountered prostate samples.

**Figure 4:**
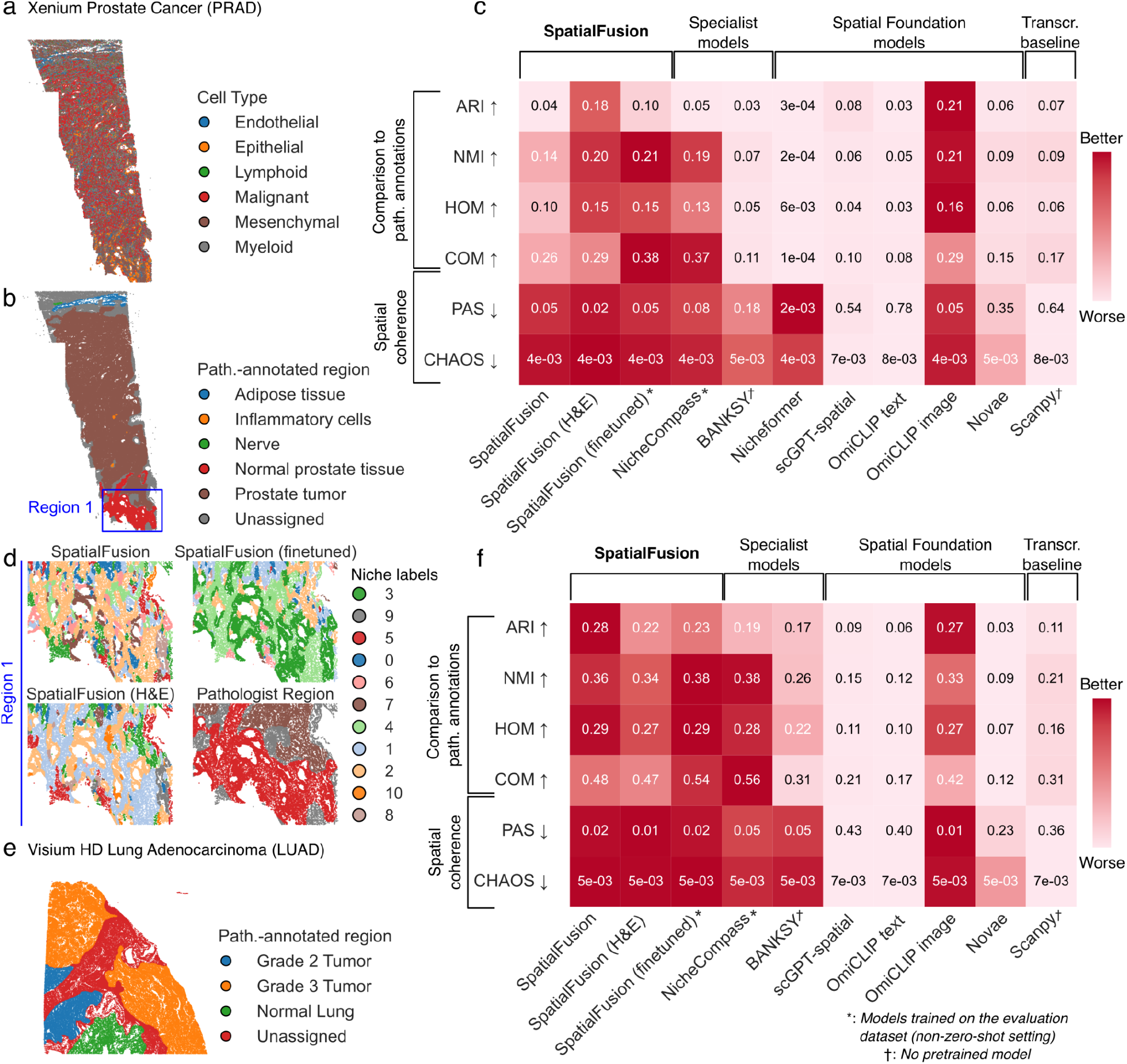
SpatialFusion performs competitively in unseen tissue (Xenium prostate cancer, PRAD) and unseen technology (Visium HD lung adenocarcinoma, LUAD). **a**, Expert-annotated cell types displayed in spatial coordinates for the PRAD sample. **b**, Pathologist-defined regions in PRAD, with one region of interest highlighted. **c**, Benchmark comparing SpatialFusion with specialist niche-detection methods, spatial foundation models, and a transcriptomic baseline, analogous to Fig. 3c. All models were applied to the PRAD sample with default parameters and embeddings were clustered using Leiden clustering (n = 11). **d**, Zoomed view of a normal prostate region showing niches identified by SpatialFusion, SpatialFusion (fine-tuned), and SpatialFusion (H&E-only), alongside pathologist annotations overlaid on the H&E image. **e**, Pathologist-defined regions in the LUAD Visium HD sample. **f**, Benchmark for the LUAD sample, equivalent to panel c, showing performance of SpatialFusion and other models on an unseen experimental technology.

Confusion matrix analysis revealed that finetuning primarily improved performance in normal prostate regions (Fig. 4d, Supplementary Fig. 4). The zero-shot model struggled to distinguish normal from malignant areas, whereas the SpatialFusion (H&E) variant performed comparably to NicheCompass and to the image-only variant of OmiCLIP (OmiCLIP-image).

We further evaluated SpatialFusion on an unseen technology by applying it to a LUAD Visium HD sample annotated with grade 2, grade 3, and normal lung regions^27^ (Fig. 4e, Supplementary Fig. 5). In this benchmark, the zero-shot SpatialFusion model outperformed all other methods in ARI, HOM, and spatial coherence metrics, while NicheCompass achieved higher NMI and COM, and OmiCLIP ranked closely behind (Fig. 4f, Supplementary Fig. 5). Remarkably, SpatialFusion recovered biologically meaningful spatial niches in this new technology without any finetuning, demonstrating strong generalization across platforms. BANKSY performed poorly in this context. The fine-tuned SpatialFusion variant further improved performance, surpassing all methods except in COM, where NicheCompass remained slightly higher, and in ARI, where OmiCLIP led. NicheFormer was not evaluated in this setting as it does not support Visium HD data. Collectively, these results highlight SpatialFusion’s robust generalization across both unseen tissues and technologies, with fine-tuning providing additional gains in novel tissue contexts while the zero-shot model performs strongly on new technological platforms.

Having established SpatialFusion’s competitive performance across diverse benchmarking scenarios, we next applied the method to illustrate its utility for biological discovery in additional, larger cohorts lacking ground-truth pathologist annotations.

### SpatialFusion uncovers a pre-malignant niche in normal-appearing mucosa adjacent to colorectal cancer

To demonstrate how SpatialFusion can enable biological discovery in real-world settings where cohorts contain multiple samples, lack ground-truth pathologist annotations, and include distinct conditions that may drive shifts in tissue organization, we next applied the method to a Visium HD colorectal cancer (CRC) cohort comprising three tumor samples (P1CRC, P2CRC, P5CRC) and two normal-adjacent tissue (NAT) samples (P3NAT, P5NAT)^28^. SpatialFusion identified 18 spatial niches across patients (Fig. 5a, Supplementary Fig. 6, Methods). Several niches were predominantly found in CRC samples (niches 0, 2, and 5), whereas others were largely restricted to NAT samples (niches 1, 7, 9, and 12) (Fig. 5b). Most niches exhibited heterogeneous cell-type composition, with the dominant cell type representing less than 60% of the niche (Fig. 5c).

**Figure 5:**
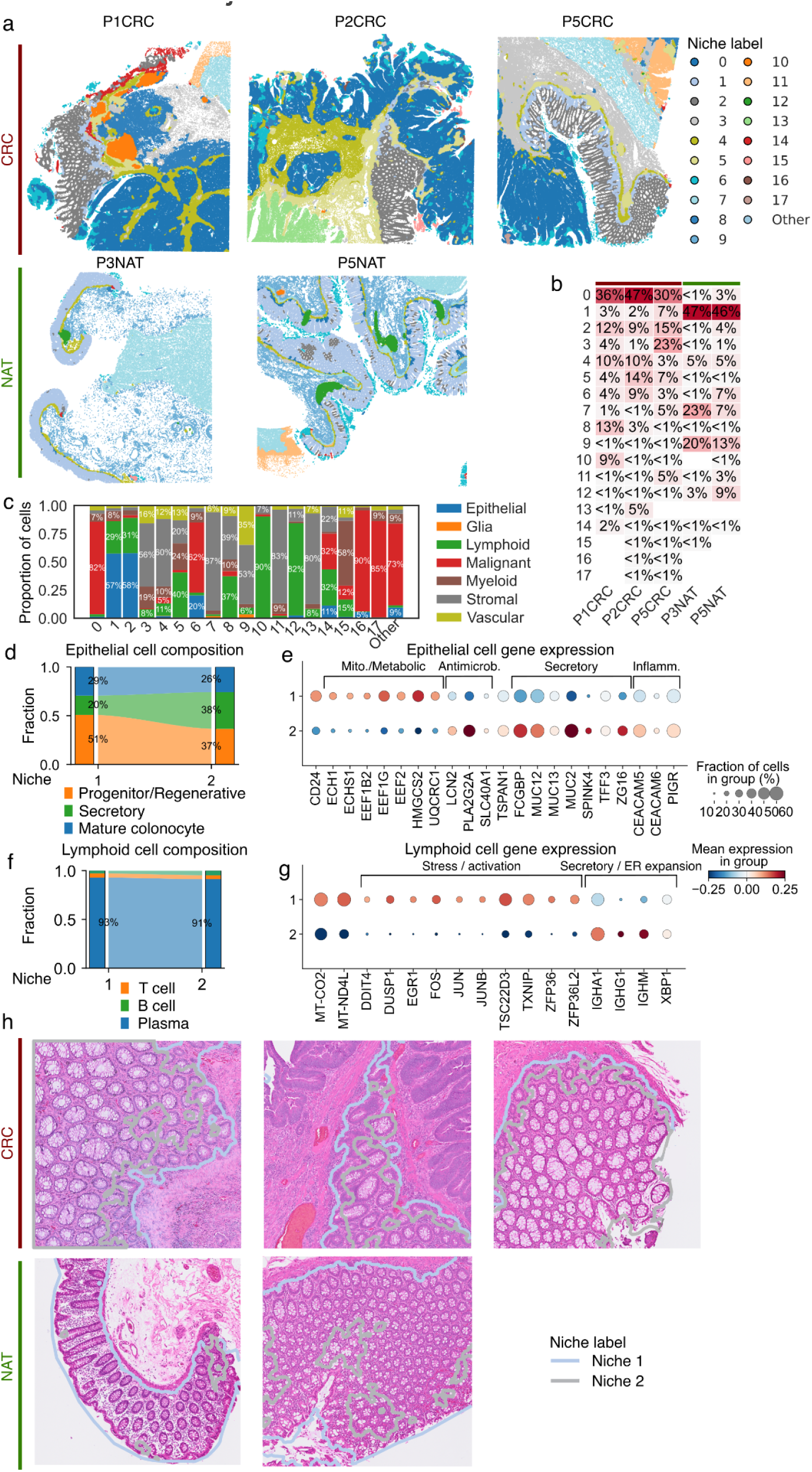
SpatialFusion identifies a pre-malignant niche within morphologically normal mucosa adjacent to colorectal tumors. **a**, Spatial coordinates of cells colored by SpatialFusion-derived niches for three colorectal cancer (CRC) samples (top row) and two normal-adjacent tissue (NAT) samples (bottom row). **b**, Distribution of niches across samples, shown as the percentage of cells in each sample assigned to each niche. **c**, Cell type composition of SpatialFusion-defined niches. **d**, Alluvial plot depicting shifts in epithelial cell subtype composition between niche 1 (homeostatic mucosa) and niche 2 (defensive mucosa). **e**, Dot plot showing Z-scores of genes upregulated in niche 1 versus niche 2 epithelial cells, grouped by functional category. **f**, Alluvial plot showing changes in lymphoid cell subtype composition between niches 1 and 2. **g**, Dot plot showing Z-scores of genes upregulated in niche 1 versus niche 2 lymphoid cells, grouped by functional category. **h**, H&E zooms of morphologically normal mucosa from CRC and NAT samples, with niche 1 and niche 2 boundaries overlaid.

We focused on niches 1 and 2, which exhibited highly similar overall cell-type compositions, dominated by epithelial and lymphoid cells, but differed markedly in the conditions in which they appeared (Fig. 5b). Both niches localized to morphologically normal-appearing mucosa; however, niche 1 was predominantly found in NAT samples, whereas niche 2 occurred almost exclusively in CRC samples, specifically in normal-appearing mucosa adjacent to tumor tissue (Supplementary Fig. 6) These findings suggest that niche 2 represents a shifted or defensive state of the mucosa near the tumor, which might be consistent with the phenomenon of field cancerization in CRC^29–31^. Notably, these regions could be identified by a pathologist in only one sample exhibiting cytomorphological features of dysplasia; however, in the other samples, such regions could not be distinguished on H&E staining, indicating that both would be classified as normal in routine diagnostic practice (M.S.). Additionally, these niches were also uncovered using the H&E only SpatialFusion embedding (Supplementary Fig. 6).

To delineate the molecular differences between the defensive state niche 2 and the homeostatic niche 1, we compared their cellular and transcriptional compositions. Niche 2 was enriched in secretory epithelial cells and depleted in progenitor or regenerative epithelial populations (Fig. 5d). Differential gene expression analysis revealed that epithelial cells in niche 2 upregulated secretory, antimicrobial, and inflammatory programs, whereas those in niche 1 expressed higher levels of mitochondrial and metabolic genes, indicating a shift from a homeostatic state in niche 1 to a defensive epithelial state in niche 2. The lymphoid compartment, composed primarily of plasma cells in both niches, also displayed a transcriptional reprogramming: plasma cells in niche 2 expressed genes associated with secretion and endoplasmic reticulum expansion, while those in niche 1 upregulated stress- and activation-related genes. Together, these observations suggest that morphologically normal mucosa adjacent to tumor tissue adopts a secretory and inflamed phenotype, mucosa that was not identified on H&E alone by a pathologist.

Histological inspection supported these molecular findings. Cells within niche 2 exhibited larger vacuoles than those in niche 1, consistent across samples (Fig. 5h). Since mucin appears as clear unstained regions in H&E images^32,33^, these vacuoles likely correspond to increased mucin production, compatible with the transcriptomic evidence of a more secretory phenotype. Finally, pathway analysis revealed elevated androgen signaling and reduced JAK–STAT activity in niche 2 across patients (Supplementary Fig. 6), suggesting that altered pathway activation may contribute to the pre-malignant reprogramming of normal-appearing mucosa near tumor tissue.

### SpatialFusion uncovers niches predictive of primary tumor stage in non-small cell lung cancer

To demonstrate the information richness of SpatialFusion embeddings, we applied the model to a Visium HD cohort of ten untreated non–small-cell lung cancers (NSCLC) generated in-house for this study, consisting of five lung adenocarcinomas (LUAD) and five squamous cell carcinomas (LUSC) spanning stages I–IV, with the goal of predicting clinical characteristics. SpatialFusion identified 18 reproducible niches across patients (Fig. 6a, Supplementary Fig. 7). Several niches were preferentially associated with one histological subtype, such as niches 6 and 14, which were enriched in LUSC and LUAD respectively, whereas others, including niches 0 and 12, were shared across both subtypes. Niches varied in composition, with some dominated by malignant cells and others composed primarily of tumor microenvironment (TME) cells or mixtures of both (Fig. 6b).

**Figure 6:**
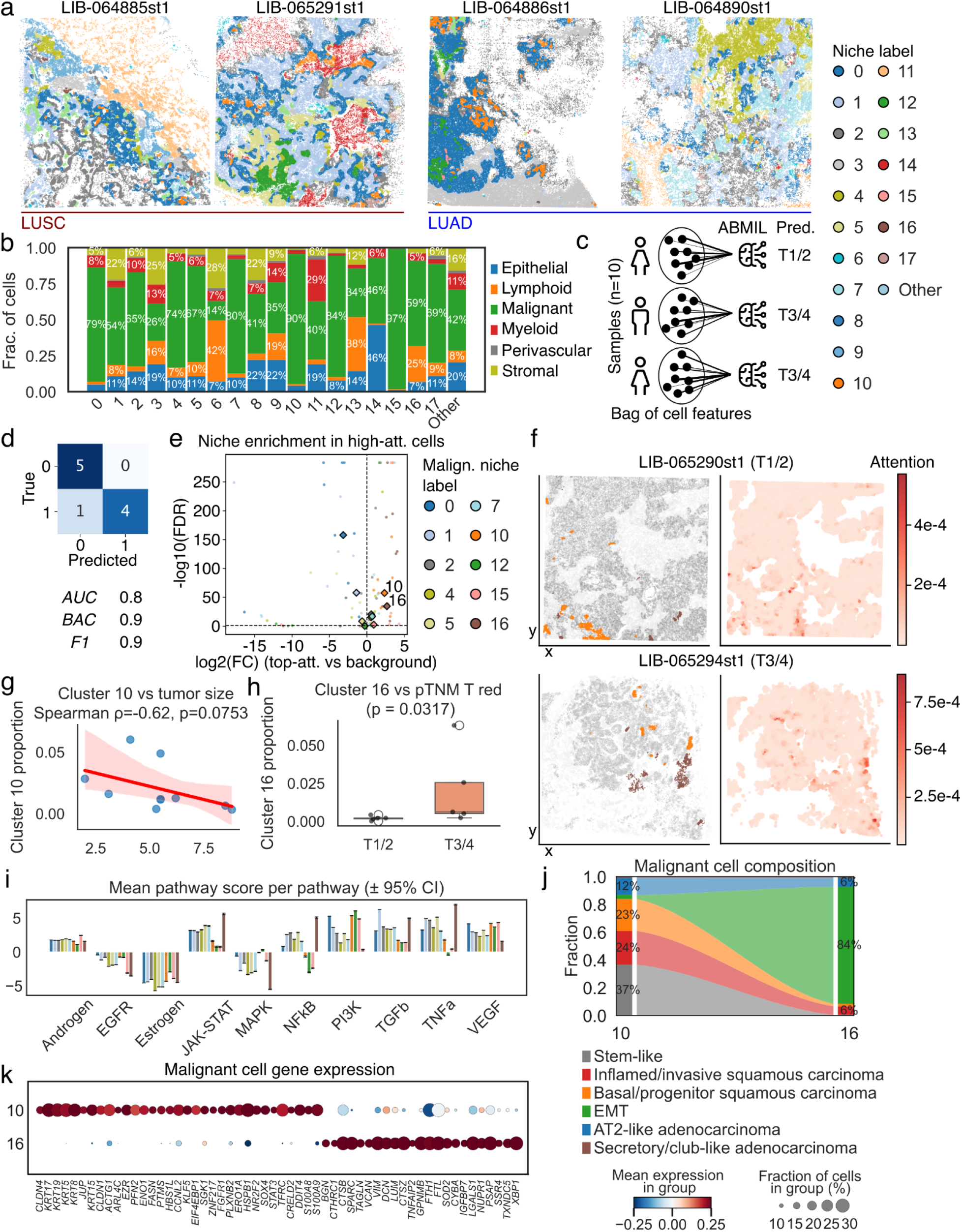
SpatialFusion identifies malignant niches predictive of pathologic tumor stage in a Visium HD non–small-cell lung cancer (NSCLC) cohort (n = 10). **a**, Representative lung squamous cell carcinoma (LUSC) and lung adenocarcinoma (LUAD) samples from the cohort, with cells colored according to SpatialFusion-derived niches. **b**, Cell type composition of SpatialFusion niches. Niches containing more than 50 percent malignant cells are designated as malignant niches. **c**, Overview of the attention-based multiple instance learning (ABMIL) framework. Bags of randomly sampled cells from each patient are used to train an ABMIL classifier to distinguish lower-stage (pT1/2) from higher-stage (pT3/4) tumors. **d**, Leave-one-out cross-validation performance of the ABMIL model, including confusion matrix and associated metrics: area under the receiver-operating characteristic curve (AUC), balanced accuracy (BAC), and F1 score. **e**, Volcano plot identifying niches enriched among the top five percent of high-attention cells. Enrichment is assessed by Fisher’s exact test, with log2 fold change on the x-axis and −log10 FDR-adjusted p-value on the y-axis. **f**, Representative pT1/2 and pT3/4 samples. Left panels show spatial distributions of niche 10 (orange), niche 16 (brown), other malignant niches (dark gray), and all remaining niches (light gray). Right panels display the learned attention map, with deeper red indicating higher attention weights. **g**, Regression analysis showing the association between niche 10 prevalence and tumor size (cm). **h**, Boxplot showing increased prevalence of niche 16 in higher-stage tumors (pT3/4). **i**, Mean PROGENy-estimated pathway activation scores for malignant niches. **j**, Alluvial plot showing shifts in malignant-cell subtype composition between niche 10 and niche 16. **k**, Dot plot showing Z-scores of genes upregulated in niche 10 and niche 16 malignant cells.

Given their potential relevance to disease progression, we focused on niches enriched in malignant cells and evaluated whether SpatialFusion embeddings could predict pTNM T stage. Multiple instance learning (MIL) is a weakly supervised approach in which labels (in this case pTNM T stage) are provided at the sample level, while individual instances (in this case cells) remain unlabeled. MIL is well suited for histopathology and spatial omics, where patient-level outcomes are known but the cellular drivers of those outcomes are not. In particular, we trained an attention-based multiple instance learning (ABMIL)^34^ model on the SpatialFusion embeddings to classify patients into early-stage (pT1/2) or late-stage (pT3/4) groups and assessed performance using leave-one-out cross-validation (Fig. 6c, Methods). The model performed well, misclassifying only one patient and achieving an AUC of 0.8, balanced accuracy of 0.9, and F1-score of 0.9 (Fig. 6d). Notably, the misclassified sample had the lowest number of cells (LIB-064887st1; n = 16,109). When trained on raw gene expression rather than SpatialFusion embeddings, the same model performed near randomly (AUC = 0.36, BAC = 0.5, F1 = 0.44), underscoring the added predictive value of multimodal spatial representations, and consistent with previous findings in NSCLC^35^.

To examine how the model made its predictions, we analyzed the attention weights learned by the ABMIL classifier. We selected the top 5 percent of cells ranked by attention score and tested whether specific niches were overrepresented among these highly weighted cells relative to their overall prevalence (Fig. 6e, Methods). Although elevated attention values appeared broadly across malignant regions, high-intensity attention signals occurred more frequently and more prominently within niches 10 and 16. Both niches were significantly enriched among top-attention cells (cluster 10: median log2FC = 2.3, χ^2^(20) = 3868.93; cluster 16: median log2FC = 2.7, χ^2^(20) = 4731.07; Fisher’s combined probability test, both p < 10^-300^; n = 10 samples), indicating that the classifier relied disproportionately on these microenvironments for stage prediction (Fig. 6e-f, Supplementary Fig. 8).

We next examined the clinical correlates of these niches. The proportion of niche 16 was significantly (Mann–Whitney U = 1, median difference = 0.0044, p = 0.032, n = 9) associated with later T stage, whereas niche 10 showed a borderline association with smaller tumor size (Spearman correlation: ρ = −0.62, 95% CI [−0.91, 0.08], t(7) = −2.09, p = 0.075, n = 9) but not with T stage directly (Fig. 6g). These patterns suggest that the ABMIL model leverages the relative abundance of niches associated positively (niche 16) and negatively (niche 10) with tumor progression. Molecular characterization revealed striking differences between the two malignant niches (Fig. 6i). Niche 16 displayed high activation of immune-related pathways, including JAK–STAT, TNFα, and NF-κB, and low activation of PI3K and VEGF signaling. Niche 10 exhibited the inverse pattern, with strong PI3K and VEGF activity and low immune-pathway activation. Notably, both niches were composed predominantly of malignant cells, which suggests that the immune-related pathway activity observed in niche 16 likely reflects tumor-intrinsic inflammatory programs rather than signaling driven by the surrounding microenvironment. This interpretation aligns with prior reports describing inflammation-activated, pro-tumor transcriptional states^36–38^, whereas niche 10 corresponds to a more canonical oncogenic signaling phenotype^39^.

We then assessed whether malignant cell phenotypes differed between niches. Malignant cells in niche 10 and niche 16 shifted from heterogeneous LUAD (AT2-like, stem-like) and LUSC (basal/progenitor, inflamed/invasive) transcriptional states toward an epithelial-to-mesenchymal transition (EMT) phenotype (Fig. 6j). This shift was observed in both LUAD and LUSC, though niche 16 in LUSC consisted almost exclusively of EMT-like malignant cells, whereas niche 16 in LUAD retained a mixture of EMT and AT2-like secretory cells (Supplementary Fig. 7). Gene expression profiles were consistent with this pattern: niche 10 malignant cells expressed keratins (*KRT5, KRT6A*), junctional genes (*CLDN1, CLDN4, JUP*), and proliferation markers (*KLF5, CCNL2*), whereas niche 16 malignant cells upregulated ECM-remodeling and mesenchymal genes (*SPARC, DCN, VIM, TAGLN*), stress-response genes (*XBP1, TXNDC5, NUPR1*), and immune-evasion markers (*LGALS1*).

Finally, niche 16 contained a substantial lymphoid compartment enriched for IgA/IgG plasma cells (Fig. 6b, Supplementary Fig. 7), consistent with a terminally differentiated state. Taken together, these findings indicate that the two niches most predictive of T stage represent distinct malignant microenvironments. Niche 10 corresponds to a basal, proliferative, well-vascularized tumor state, whereas niche 16 reflects a more aggressive, inflammation-activated malignant program marked by EMT, tumor-intrinsic immune-pathway activation, and features associated with immune evasion. These results show that SpatialFusion identifies in an unsupervised manner malignant niches associated with tumor progression, each defined by distinct molecular profiles and marked differences in pathway activation, including pathways with FDA-approved targeted therapies such as JAK–STAT. This emphasizes the translational potential of the approach. Notably, the zero-shot SpatialFusion embeddings predicted pathological tumor stage in a very small cohort (n = 10) despite the absence of stage supervision, underscoring the informative nature of the learned spatial representations.

## Discussion

We introduce SpatialFusion, a lightweight multimodal deep learning framework that integrates spatial transcriptomics and histopathology to identify biologically meaningful spatial niches. The method has three main advantages. First, its multimodal design enables application to paired ST–H&E datasets as well as H&E-only cohorts at a (inferred) single-cell resolution, allowing broad use across experimental settings and supporting the discovery of niches defined by both transcriptional and morphological features. Second, SpatialFusion explicitly incorporates pathway-level supervision to distinguish spatial neighborhoods based on molecular activation states rather than cell identity alone. Third, by relying on frozen foundation models, the framework is computationally efficient and can be trained on a standard GPU-equipped laptop, with fewer than 300,000 trainable parameters and low VRAM requirements (<5 MB for training and <2 MB for inference).

We benchmark SpatialFusion against specialist niche-detection approaches (NicheCompass^9^ and BANKSY^10^) and recent spatial foundation models (OmiCLIP^15^, scGPT-spatial^12^, Nicheformer^11^, Novae^14^). Across tissue types and data modalities, SpatialFusion and its H&E-only variant perform competitively, including in settings involving unseen tissues and unseen technologies such as Visium HD. The method also reliably identifies fine-grained niches with distinct pathway-activation signatures, which are often missed by unimodal or single-cell–centric architectures.

To evaluate biological utility, we applied SpatialFusion to two additional Visium HD cohorts that were not used in training. In CRC, the model identified two niches with similar overall cell-type composition but divergent spatial distributions: niche 1 was enriched in uninvolved normal-adjacent tissue, while niche 2 appeared exclusively in morphologically normal mucosa proximal to tumor. Although indistinguishable on H&E, niche 2 displayed a shift in epithelial and plasma cell phenotype, including increased secretory, antimicrobial, and terminal differentiation programs. These observations suggest that niche 2 represents a defensive or stress-associated state. The altered epithelial activity and accompanying B-cell maturation raise the possibility that tumor-adjacent mucosa experiences paracrine or microenvironmental influences that promote immune modulation and matrix remodeling. This interpretation is consistent with the concept of field cancerization in CRC^29,31^. Prior spatial studies in CRC have largely focused on intratumoral niches^28,40–42^, with few examining normal-appearing mucosa; the original cohort analysis did report goblet-cell changes near the tumor boundary^28^, consistent with our results. Importantly, a board-certified pathologist (M.S.) confirmed that these mucosal regions would be classified as normal in routine diagnostic practice, underscoring the complementary value and potential utility of SpatialFusion.

SpatialFusion also revealed clinically relevant spatial organization changes in NSCLC. Multimodal embeddings supported accurate prediction of tumor stage using a weakly supervised ABMIL model, outperforming a gene expression–only baseline. Two malignant niches contributed disproportionately to model decisions: niche 10, associated with smaller tumors, showed a proliferative and well-vascularized profile, whereas niche 16 exhibited an invasive phenotype marked by elevated JAK–STAT, TNFα, and NF-κB activity and was associated with higher T stage. These findings suggest that distinct malignant microenvironments encode architectural and molecular correlates of local tumor progression. While prior spatial analyses in NSCLC have linked microenvironmental structure to immune state and treatment response^35,43–47^, to our knowledge, discrete spatial niches associated with T stage have not been previously described. Together, these findings illustrate SpatialFusion’s capacity to identify biologically meaningful niches in a fully unsupervised manner, revealing microenvironmental architectures that track with tumor progression and may support future prognostic or therapeutic stratification efforts.

There are several limitations to this study. First, benchmarking relies on pathologist annotations as a proxy for ground truth, however, these labels are coarse-grained, based solely on H&E morphology, and fail to capture molecular heterogeneity. Although we assess molecular coherence and use metrics such as homogeneity and completeness to account for fine-grained structure, this limitation remains. Second, transcript diffusion and assignment based on segmentation methods in Visium HD data affects cell type assignment, and despite multiple rounds of annotation, some contaminating transcripts persist; future work could incorporate explicit decontamination. Third, SpatialFusion depends on frozen scGPT and UNI embeddings, which makes the model lightweight but also propagates any biases present in these foundation models. Fourth, we do not systematically evaluate sensitivity to architectural choices such as graph construction or pathway sets. While the PROGENy pathways used here are broadly relevant across tissues and applicable beyond oncology (e.g., fibrosis or neurodegeneration), we focused on pathways with strong relevance to cancer biology (those with approved targeted therapies, under clinical investigation, or indirectly targetable) potentially biasing the latent space toward cancer-specific signaling. Fifth, there is no definitive ground truth for the number of spatial niches. Although ovarian cancer benchmarks show robustness across clustering resolutions, the numbers selected for CRC and NSCLC remain somewhat arbitrary. Finally, the additional cohorts are modest in size, and both the niche–stage associations and the biological identity of inferred niches require validation in larger datasets and with orthogonal experimental assays. The results should therefore be interpreted as hypothesis-generating.

In summary, SpatialFusion provides a lightweight and versatile framework for multimodal spatial analysis, capable of uncovering morpho-molecular niches with functional and clinical relevance. By integrating histopathology, transcriptomics, and pathway-level signals, the method uncovers tissue niches defined by distinct molecular activation patterns that single modalities cannot resolve. Our results across CRC and NSCLC demonstrate that SpatialFusion can identify previously unrecognized niches, including defensive states in normal-appearing mucosa and malignant niches associated with tumor invasiveness. As spatial profiling technologies continue to expand across tissue types and clinically relevant contexts, methods such as SpatialFusion offer a scalable and interpretable foundation for characterizing disease-associated tissue states and generating mechanistic hypotheses for future experimental validation.

## Methods

### Datasets used in the study

SpatialFusion was trained on a subset of Xenium spatial transcriptomics (ST) experiments from the HEST1-k dataset^16^. We used 51 samples for training and held out 8 samples for testing (sample IDs provided in Supplementary Table S2). The HEST1-k data can be downloaded following the instructions at: https://huggingface.co/datasets/MahmoodLab/hest.

For benchmarking on Xenium data, we used two held-out HEST1-k test samples, NCBI875 (https://www.10xgenomics.com/products/xenium-in-situ/preview-dataset-human-breast, in situ sample 1, replicate 1) and TENX157 (https://www.10xgenomics.com/datasets/xenium-prime-ffpe-human-prostate), as well as an ovarian cancer (OVCA) Xenium sample not included in HEST1-k (https://www.10xgenomics.com/datasets/xenium-prime-ffpe-human-ovarian-cancer). For Visium HD benchmarking, we used data from a 10X publicly released sample (https://www.10xgenomics.com/datasets/visium-hd-cytassist-gene-expression-human-lung-cancer-post-xenium-expt, Experiment 1, HD only).

Pathologist annotations for the Xenium OVCA, Xenium BRCA, and Visium HD LUAD datasets were obtained by downloading the annotated images and retracing them in QuPath^48^. For OVCA, the annotated image was obtained directly from the 10x Genomics website; for BRCA^49^, we used the image provided in the associated *Nature Communications* publication; and for LUAD, we reconstructed annotations based on this recent study^27^. For the prostate cancer (PRAD) Xenium sample, a board-certified pathologist (M.S.) generated annotations in QuPath. All cells within annotated regions were assigned the corresponding pathologist-defined niche.

For downstream analyses, we first used the five publicly available Visium HD colorectal cancer (CRC) and normal-adjacent tissue (NAT) samples (https://www.10xgenomics.com/platforms/visium/product-family/dataset-human-crc) from the recent publication^28^. We additionally used ten non-small-cell lung cancer (NSCLC) samples from an in-house Visium HD cohort (will be uploaded to a public repository upon acceptance of the manuscript).

### Overview of SpatialFusion methodology

The architecture consists of three main modules: a unimodal encoding module, a multimodal alignment module, and a spatial encoding module.

#### Unimodal encoding module

In this module, each cell is represented by its raw gene expression counts and a 256 × 256–pixel H&E patch centered on the cell. Most histology images have a resolution of ~0.2 microns per pixel (MPP), as reported in the HEST1-k metadata or in the Xenium and Visium HD datasets, corresponding to physical patches of approximately 50 × 50 µm.

For the transcriptomic modality, we compute scGPT^1^ embeddings by applying the whole-human pretrained scGPT model with default parameters, using adapted code from https://github.com/microsoft/zero-shot-scfoundation.

For the imaging modality, we compute UNI2^3^ embeddings by applying the UNI model to each H&E patch; patches are zero-padded when needed to maintain a consistent 256 × 256 input size. Image-to-transcript coordinate alignment depends on the dataset:

- For HEST1-k, H&E images are already aligned to transcript coordinates by the original authors.
- For the OVCA Xenium sample, we use the alignment file provided by 10x Genomics to convert IF-based coordinates into high-resolution H&E coordinates.
- For Visium HD, spatial coordinates are natively defined in the high-resolution image space.

No parameters are trained in this module; only forward inference through the pretrained scGPT and UNI models is performed. The output of this module is a lower-dimensional embedding 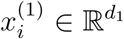 with *d*_1_ = 1536 the output size of UNI2 and 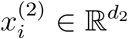 with *d*_2_ = 512 the output size of scGPT.

#### Multimodal alignment module

For each cell *i* in the dataset, we obtain an aligned pair of modality-specific features 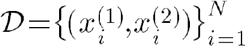.

These features are standardized per sample before being passed to the encoders.

To align the two modalities, we construct a multimodal autoencoder (mAE) composed of two independent encoders and two corresponding decoders. Each modality is mapped to a shared latent space of dimension *L* = 64.

The encoders are parameterized functions,

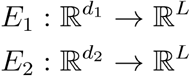

implemented as multilayer perceptrons (MLPs) with a single hidden layer of size 64 and ReLU activations.

For each cell *i*, we thus obtain the lower dimensional representation:

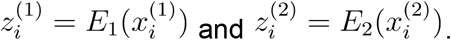

Two decoders, mirroring the encoder architecture, project latent vectors back into their respective modality spaces:

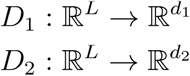

These decoders enable both within-modality reconstructions:

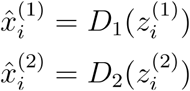

and cross-modality reconstructions:

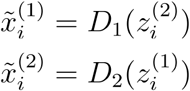

The model is trained using the total loss:

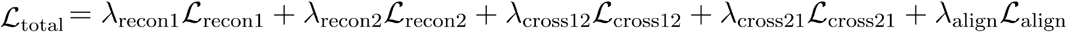

This objective includes:

1. Within-modality reconstruction losses to enforce accurate self-reconstruction:

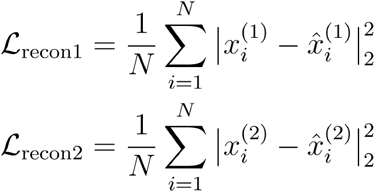
2. Cross-modality reconstruction losses to promote latent representations that are sufficiently informative to reconstruct the opposite modality, we use:

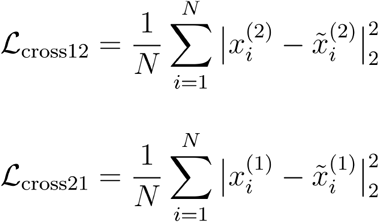
3. Latent alignment penalty to encourage consistent embeddings for paired modalities:

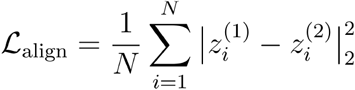

The weights used in this work were:

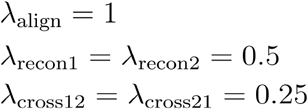

Weights were selected to prioritize accurate within-modality reconstruction over cross-modality reconstruction, while assigning a relatively higher weight to the latent alignment term to encourage consistency between modalities.

Parameters of all four networks Θ = *E*_1_, *E*_2_, *D*_1_, *D*_2_ were optimized using stochastic gradient descent with the Adam optimizer. The objective was minimized over 25 epochs using minibatches of size 128, a learning rate of (10^−3^), and GPU acceleration, which are standard choices.

For downstream analyses, latent embeddings were computed in inference mode:

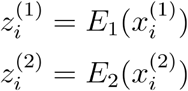

In ablation experiments, we compare several embedding strategies:

- the joint embedding 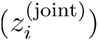 obtained by averaging 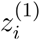 and 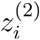;
- concatenated embeddings;
- unimodal embeddings, using 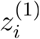 (H&E only embedding) or 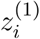 (RNA-only embedding).

We also evaluate an ablated mAE trained without cross-modality losses, i.e. using the truncated objective

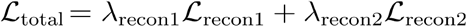

referred to as the mAE without cross-modality.

#### Spatial encoding module

We then train a graph convolutional masked autoencoder (GCMAE) to obtain latent neighborhood representations. For each sample, spatial coordinates associated with each single-cell were used to construct a k-nearest-neighbor (kNN) graph, *k* = 30. Let the set of nodes be

*V* = 1,..., *N* each with coordinates *c*_*i*_

A spatial adjacency graph was defined as:

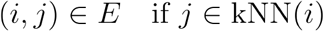

resulting in an undirected graph

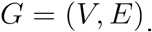

Each node was associated with a feature vector derived from joint embeddings of two modalities (e.g., from the paired autoencoder described above):

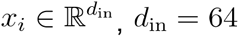 in our experiments.

We standardized the features across all cells of a single slide before graph construction.

To enforce robustness and encourage the GCN to reconstruct missing information, a randomly sampled subset of nodes was masked during training. Let

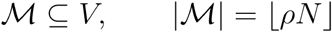

where *ρ* is the node masking ratio. We choose *ρ* = 0.9 for our experiments. This high masking ratio was motivated by experimental results from the GraphMAE work showing better performance with high masking ratios^50^.

A learnable mask token

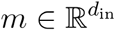

replaced the input features for nodes in ℳ:

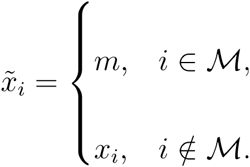

Gaussian noise was added to all node features:

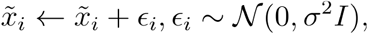

followed by dropout with probability *p*. Gaussian perturbations encourage the model to learn smooth, noise-invariant representations rather than overfitting sample-specific variation and act as a regularizer^51,52^. Dropout reduces feature co-adaptation and improves robustness in graph neural networks^50,53^. Together, these regularizers promote stable neighborhood embeddings and improve generalization.

The GCMAE encoder consisted of (*L*) sequential graph convolutional layers (*L* = 2 in our experiments). Because we construct a 30–nearest-neighbor graph, the theoretical receptive field after *L* layers includes all nodes reachable within 2 hops, which could be as large as 30^2^ = 900) *nodesifthegrap* were unconstrained. However, this upper bound substantially overestimates the *effective* receptive field in spatial transcriptomics data.

Thus, although the theoretical receptive field is at most 900 nodes, geometric constraints imposed by the underlying 2D tissue architecture limit the effective receptive field to approximately 120 cells in practice. This estimate is consistent with empirically observed neighborhood densities in spatial transcriptomics datasets.

The encoder consists of *L* sequential graph convolutional layers (*L* = 2 in our experiments). Because we construct a 30–nearest-neighbor graph, the theoretical receptive field after *L* layers includes all nodes reachable within 2 hops, which could be as large as 30^2^ = 900 nodes if the graph were unconstrained. However In practice, the spatial graph is embedded in a 2-dimensional tissue plane, where cells form an approximately locally uniform point cloud. Under this assumption, the number of nodes within *h* hops grows with the area of the neighborhood rather than exponentially. If the 30 nearest neighbors of a cell lie within radius *r*, then the 2-hop neighborhood corresponds to nodes within radius 2*r*. Because area in 2D scales as (2*r*)^2^ = 4*r*^2^, the effective receptive field after 2 GCN layers is approximately 2^2^ × *k* = 4*k* where *k* = 30. This yields an effective receptive field of roughly: 4 × 30 = 120 cells.

We implement the GCN with standard parameters and layers. For layer *l*:

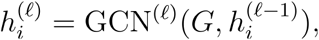

where 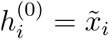.

Each layer applied a DGL GraphConv operator:

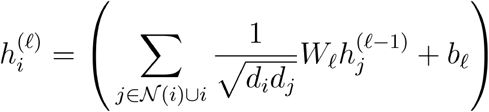

where *d*_*i*_ is the degree of node *i*.

Layer normalization was applied after each layer:

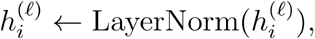

and for all but the final layer:

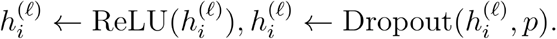

The final latent representation was:

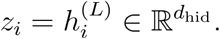

where *d*_hid_ = 10 in our experiments. We then used this latent representation for two tasks. The first is reconstruction only on the masked nodes. A two-layer MLP decoder mapped latent representations back to the input feature space:

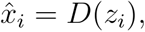

where

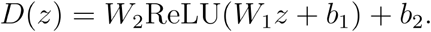

Reconstruction was computed only on masked nodes:

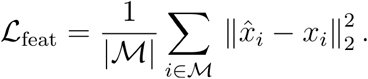

The second supervised task involved predicting pathway activation levels. For each cell, we estimated pathway scores using PROGENy^19^, following the workflow described in their spatial transcriptomics tutorial (https://decoupler.readthedocs.io/en/latest/notebooks/spatial/rna_visium.html). Briefly, we first applied spatial weighting to impute gene expression in a spatially informed manner, then computed PROGENy pathway activities on these spatially smoothed profiles.

We selected ten PROGENy signaling pathways with established relevance to cancer biology. These included pathways with approved targeted therapies (EGFR, androgen, estrogen, JAK–STAT, VEGF, MAPK, PI3K), those currently under clinical investigation (TGFβ), and additional pathways that can be indirectly modulated (NFκB and TGFA). For each cell *i* this procedure generated a vector of pathway activation estimates, which we used as regression targets and denote as 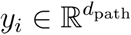.

An additional regression head predicted pathway scores:

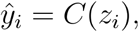

where *C* is a linear classifier:

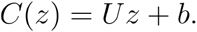

A Huber loss (Smooth L1 loss) quantified prediction error:

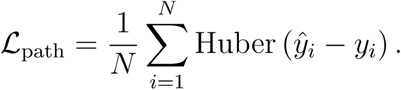

Finally, to prevent the latent space from collapsing, we use the standard addition of an *l*_*2*_ regularization term applied to the representations:

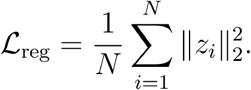

The model was trained by minimizing the weighted combination of reconstruction loss on masked nodes, pathway regression loss, and a latent regularization term. The full objective was:

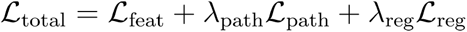

In this study, the weights were chosen to prioritize feature reconstruction over pathway regression, with a modest regularization term added to prevent it from dominating the total loss:

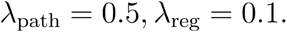

The GCNAutoencoder parameters

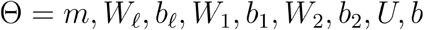

were optimized using Adam with learning rate *η* = 10^−3^. Training was performed in a standard fashion for 25 epochs with batch size 1. All training was executed on a CUDA-enabled GPU.

To enable scalable training on large spatial graphs, we partitioned each tissue graph into a collection of overlapping local subgraphs using spatial clustering. Let *G* = (*V,E*) be the full graph with | *V* | = *N* nodes and associated spatial coordinates *c*_*i*_ ∈ ℝ^2^. Rather than using a fixed sliding window, we employed a spatially informed clustering strategy to select local neighborhoods.

First, we defined a target stride parameter *T*, which determines the density of cluster centers across the tissue. The number of cluster centers was computed as

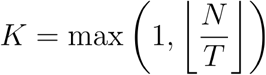

and *K* spatial clusters were identified by MiniBatch (k)-means applied to the coordinate matrix *C* = [*c*_1_, …, *c*_*N*_]^T^. For each cluster *k* ∈ 1, …, K, we collected the set of nodes assigned to that cluster:

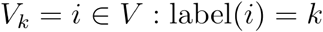

To maintain computational tractability, a maximum subgraph size *S* was imposed. If |*V*_*k*_| > *S*, a random subset of |*V*_*k*_| = *S* nodes were sampled uniformly. Each subgraph was then constructed as the node-induced subgraph

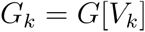

preserving all edges among the selected nodes. Node features were inherited directly from the full graph.

Setting *T* = 2500 yields roughly *N*/2500 cluster centers per slide, which provides dense spatial coverage and overlap between subgraphs while keeping the number of subgraphs manageable. Setting *S* = 5000 ensures that each subgraph contains many times the typical receptive field size, so that most nodes are far from subgraph boundaries and their 2-hop neighborhoods are largely preserved within the same subgraph. From a computational standpoint, capping subgraphs at 5000 nodes bounds memory and runtime: with 64-dimensional node features and ~30 edges per node, each subgraph remains feasible to process in mini-batches on commodity GPUs, while avoiding the quadratic costs associated with much larger induced neighborhoods.

This procedure generated a collection of spatially coherent subgraphs that sampled the tissue space and served as independent training inputs for the graph convolutional autoencoder, enabling efficient learning while preserving local spatial structure.

In our ablation study, we compare the GCMAE with and without the pathway regression loss *L*_path_.

### Baseline autoencoder

To assess the benefit of foundation-model initialization and multimodal alignment in SpatialFusion, we implemented a baseline multimodal autoencoder that receives the same inputs as the multimodal AE: a 256×256 H&E image patch and a raw gene expression (GEX) vector for each cell. Just like for the original mAE, images are resized or zero-padded to 224×224 pixels.

For each cell *i*, the inputs are:

- an image tensor 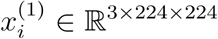, and
- a gene expression vector 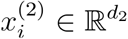.

The GEX features correspond to a soft union of genes appearing in at least 50% of training samples, standardized per sample.

Thus, each cell is represented as a paired feature:

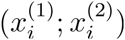

The baseline autoencoder consists of two independent encoders projecting each modality into a shared 64-dimensional latent space, and two decoders reconstructing their respective modalities. In particular, we use a ResNet18 pretrained on ImageNet, with the classification head removed, with an output feature map flattened and passed through a linear layer:

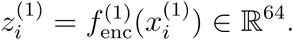

The ResNet backbone is entirely frozen, so only the projection layer is trainable.

The image decoder 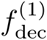 maps a latent vector *z* ∈ ℝ^*L*^ (with *L* = 64) back to an RGB image 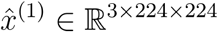. It consists of a fully connected expansion to a 3D feature tensor of shape 512 × 7 × 7, followed by five transposed convolution blocks that progressively upsample the spatial resolution:

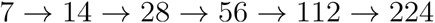

We obtain the reconstruction:

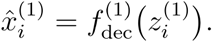

A simple, two-layer multilayer perceptron maps gene expression into the latent space:

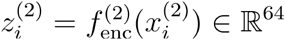

A mirrored MLP reconstructs the GEX vector:

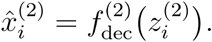

The model also produces cross-modal reconstructions:

- image → gene expression:

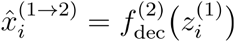
- gene expression → image:

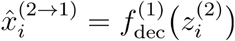

We use the same training procedure and training objective as the mAE to train the baseline AE.

### Preprocessing of HEST1-k cohort

Because cell types were not provided for the HEST1-k cohort, we generated manual annotations for all samples used in this study. Raw gene counts were first normalized using counts-per-10k (CP10k) followed by a log1p transformation. We then performed principal component analysis (PCA), constructed a k-nearest-neighbor graph in the PCA space, and applied Leiden clustering. Cell types were assigned to clusters based on canonical marker genes identified from differential expression analysis. Annotation was performed iteratively with expert review, assisted by ChatGPT^54^. The cell types were used in the ablation study to evaluate the mAE. The final cell type annotations are available on Zenodo at : 10.5281/zenodo.18377525.

### Methods used in benchmarking

To benchmark our method, we compare against two specialist models and three spatial foundation models, as well as one transcriptomic baseline.

The specialist models are:

- NicheCompass^9^: NicheCompass constructs a spatial cell–cell interaction graph and applies a graph variational autoencoder to learn embeddings that capture communication programs and local microenvironmental structure. We applied NicheCompass using default parameters following the official tutorial (https://nichecompass.readthedocs.io/en/latest/tutorials/notebooks/mouse_cns_single_sample.html).
- BANKSY^10^: BANKSY embeds each cell in a “product space” combining its own transcriptomic profile with the average profile of its spatial neighbors, followed by clustering in this augmented space. We applied BANKSY with default settings as described in the tutorial (https://github.com/prabhakarlab/Banksy_py/blob/main/slideseqv1_analysis.ipynb).

Following the authors’ recommendations for niche-discovery tasks, we used λ = 0.8.

The spatial foundation models are:

- scGPT-spatial^12^: scGPT-spatial is a spatial-omics foundation model that adapts scGPT for spatial transcriptomics. It is continually pretrained on a large multimodal corpus spanning Visium, Visium HD, Xenium, MERFISH, and related technologies. The model incorporates a neighborhood-aware reconstruction objective that conditions each cell’s reconstruction on its spatial neighbors. We applied scGPT-spatial using default parameters following the official tutorial (https://github.com/bowang-lab/scGPT-spatial/blob/main/tutorials/Multi_Modal_Integration_Demo.ipynb) and used the released pretrained weights (https://figshare.com/articles/software/scGPT-spatial_V1_Model_Weights/28356068?file=52163879).
- Nicheformer^11^: NicheFormer is a transformer-based foundation model trained on a corpus of >110 million cells spanning dissociated single-cell and spatial transcriptomics data across tissues and species. Gene expression and experimental metadata (e.g., species, assay) are tokenized and processed by a transformer to generate unified cellular embeddings. Although trained on spatial datasets, Nicheformer does not explicitly encode spatial adjacency or neighborhood structure. We applied the model with default settings following the provided tutorial (https://github.com/theislab/nicheformer/blob/main/notebooks/tokenization/xenium_human_colon.ipynb) and used the official pretrained weights (https://data.mendeley.com/preview/87gm9hrgm8?a=d95a6dde-e054-4245-a7eb-0522d6ea7dff).
- OmiCLIP^15^: OmiCLIP is a multimodal foundation model trained on ~2.2 million paired H&E patches and Visium V1/V2 spatial transcriptomics profiles. Transcriptomic profiles are converted into textual sequences (top-expressed gene symbols), while image patches are encoded using a visual backbone; contrastive learning aligns the two modalities in a shared space. Although trained on spatial datasets, OmiCLIP does not explicitly encode spatial adjacency or neighborhood structure. The model outputs separate text and image embeddings (“OmiCLIP (text)” and “OmiCLIP (image)”). We applied OmiCLIP using default parameters (https://guangyuwanglab2021.github.io/Loki/notebooks/basic_usage.html) and the publicly released pretrained weights (https://huggingface.co/WangGuangyuLab/Loki).
- Novae^14^: Novae is a graph-based spatial foundation model trained on approximately 30 million cells across multiple tissues. It uses a self-supervised graph neural network to learn spatially informed cell embeddings by integrating gene expression with spatial adjacency. The model is designed to support a range of zero-shot spatial tasks, including domain identification and batch-robust representation learning. While Novae operates natively on transcriptomic data only, the authors demonstrate a downstream application in which Novae embeddings are combined with H&E-derived image embeddings to enable multimodal analysis; multimodality, however, is not intrinsic to the model architecture. We applied Novae with default parameters following the official tutorial (https://mics-lab.github.io/novae/tutorials/main_usage/). As recommended by the authors, we used Novae’s built-in domain detection rather than applying Leiden clustering to the learned embeddings.

As a transcriptomic-only baseline, we normalize gene expression using CP10k followed by log1p transformation and apply Leiden clustering to the resulting matrix to derive clusters.

### Benchmark metrics

We applied SpatialFusion and all comparison methods to each benchmarking sample (ovarian cancer, breast cancer, prostate cancer, and Visium HD lung adenocarcinoma) to obtain low-dimensional embeddings for every cell. Using these embeddings, we constructed a k-nearest-neighbor graph and performed Leiden clustering with Scanpy. To ensure comparability across methods, we fixed the number of clusters and merged clusters containing fewer than 1% of all cells, resulting in an identical final cluster count for all approaches. These clusters are treated as spatial niches. As an exception, for Novae we used the built-in domain detection procedure recommended by the authors, rather than applying Leiden clustering to the embeddings.

Performance was assessed using the SDMBench^21^ evaluation framework. We computed four supervised metrics that compare niche assignments to pathologist annotations (adjusted Rand index (ARI), normalized mutual information (NMI), homogeneity (HOM), and completeness (COM)) as well as two spatial coherence metrics that quantify spatial smoothness of the niche labels (percentage of abnormal spots (PAS) and spatial chaos (CHAOS)). Definitions and mathematical formulations of these metrics are provided in the original publication^21^.

### Preprocessing and analysis of colorectal cancer (CRC) cohort

We applied bin2cell^55^ to the CRC Visium HD cohort to obtain single-cell–level segmentations from high-resolution H&E images, following the authors’ tutorial. Genes detected in fewer than three bins and bins with zero counts were removed prior to segmentation. Nuclei were identified using the H&E Stardist^56^ model (probability threshold = 0.01), after which nucleus masks were expanded to approximate cell boundaries. Transcripts falling within each expanded region were assigned to the corresponding cell, and cells with fewer than 30 detected transcripts were discarded.

Cell type annotation proceeded in two steps. First, within each sample, we applied the same procedure as for the HEST1-k dataset to assign broad lineages (malignant, normal epithelial, lymphoid, myeloid, stromal, perivascular) based on canonical marker gene expression. Second, for each lineage, cells from all patients were pooled and batch-corrected using BBKNN^57^. We then refined annotations by removing contaminated or mixed-identity clusters and assigning cell subtypes using lineage-specific markers. Low-complexity cells, potential doublets, and ambiguous profiles were labeled as noise and excluded from downstream analyses. PROGENy-estimated pathway scores were computed using the same procedure as for the HEST1-k training dataset. The final cell type annotations and scores for the cohort are available on Zenodo at : 10.5281/zenodo.18377525.

SpatialFusion was run separately on each sample, after which the resulting embeddings were combined into a single dataset for cohort-level analysis. Leiden clustering on this pooled embedding matrix produced 18 clusters. All clusters with fewer than 1,000 cells were grouped together and designated as “Other.”

To compare the two niches of interest, niche 1 (mucosa in normal-adjacent tissue) and niche 2 (tumor-proximal normal-appearing mucosa), we restricted analysis to epithelial or lymphoid cells belonging to these niches. Genes with mean expression below 0.5 were excluded. Differential expression between niche 1 and niche 2 cells was assessed using a Wilcoxon rank-sum test. For visualization, we displayed selected genes from the top 25 differentially expressed genes with log2 fold change > 0 in each niche. Dot plot values represent Z-scored expressions computed within the subset of analyzed cells.

### Preprocessing and analysis of non-small cell lung cancer (NSCLC) cohort

For the NSCLC cohort, we used SpaceRanger v4.0 with the high-resolution H&E image to obtain cell segmentations, which were used directly. Cell type annotation, refinement, and PROGENy pathway estimation were performed following the same procedure as for the CRC cohort, and SpatialFusion was applied to the full set of samples. For each resulting niche, we quantified cell type composition; niches containing more than 50% malignant cells were designated as malignant niches. Associated data can be found at TBD.

#### Attention-based multiple-instance learning procedure

To evaluate the predictive value of the SpatialFusion embeddings, we treated each patient as a multiple-instance learning (MIL) “bag,” where the patient-level label (pTNM T stage) is known but individual cell labels are not. Of note, in this case we only have one slide per patient. We applied an attention-based MIL (ABMIL) framework^34^, using only embeddings from malignant-niche cells, to classify patients into lower (pT1/2) versus higher (pT3/4) T stage.

Each patient *p* is represented by a bag:

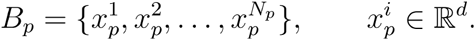

To remove inter-patient scaling effects, each bag is standardized independently:

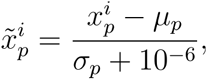

where *µ*_*p*_ and *σ*_*p*_ are the mean and standard deviation of *B*_*p*_.

During training, we randomly sub-sampled a fixed number of cells from each patient (with replacement when fewer cells were available), repeated K times per patient to generate multiple noisy bag instances per epoch.

We draw *n*_sample_ = 1,000 cells uniformly: 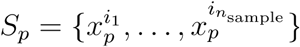.

Gaussian noise is added for robustness:

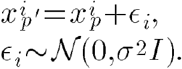

Each patient contributes *K* = 10 such noisy bags per epoch.

Each cell embedding is then scored using a nonlinear attention network:

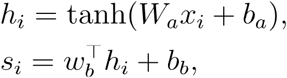

where *s*_*i*_ is the unnormalized attention score, and *W*_*a*_ ∈ ℝ ^128 × 10^, *b*_*a*_ ∈ ℝ^128^, *W*_*b*_ ∈ ℝ ^1× 128^, *b*_*b*_ ∈ ℝ^1^.

Attention weights across the bag are obtained via:

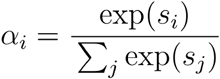

The patient-level representation is a weighted average:

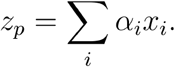

A classifier maps the pooled representation to a logit:

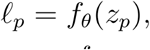

where *f* _*θ*_ is a multilayer perceptron:

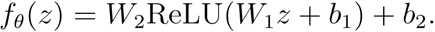

where *W*_1_ ∈ ℝ ^64 × 10^, *b*_1_ ∈ ℝ^64^, *W*_2_ ∈ ℝ ^1 × 64^, *b*_2_ ∈ ℝ^1^

The predicted probability is:

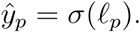

To address class imbalance, we use a positive-class weighting factor:

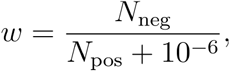

and optimize:

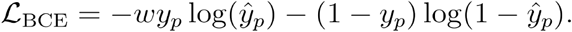

We trained an ensemble of 15 ABMIL models using leave-one-out cross-validation. For each fold, one patient was held out, and the remaining patients were used to train *R* = 15 models for 15 epochs with a learning rate of *lr* = 10^−3^

For evaluation, each model *l* in the ensemble processes the full bag for each patient *p*:

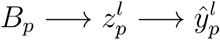 producing a complete attention map 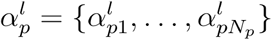.

Patient-level predictions were obtained by averaging the predicted probabilities across the ensemble. Final attention maps were computed by averaging cell-level attention weights across all models:

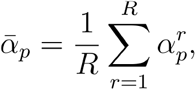

yielding stable, consensus attention profiles per patient.

To benchmark predictive performance, we trained a comparison model using the same ABMIL architecture but replacing SpatialFusion embeddings with PCA embeddings derived from log1p–CP10k–normalized gene expression for each patient.

#### Identification of niches associated with pTNM T stage

After training the ABMIL models, we assessed whether specific SpatialFusion-derived niches were disproportionately utilized by the classifier. For each patient *p*, we selected the top 5% of cells ranked by attention scores using 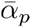. For each niche *c*, we computed its frequency among these high-attention cells, the patient 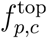 and compared it to its overall frequency in the patient’s sample, 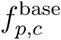. Enrichment was quantified as:

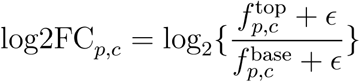

Statistical significance of enrichment was assessed using a Fisher exact test per niche–patient pair, followed by FDR correction across niches. For each niche, we report the median log2FC_*c*_ fold-change and the corresponding FDR-adjusted p-value across patients.

We next assessed clinical associations by testing the relationship between niche prevalence and pTNM T stage using a Mann–Whitney U test, and its association with tumor size (cm) using Spearman correlation.

#### Association with malignant cell phenotypic change

To investigate malignant cell state differences between niches, we restricted the analysis to malignant cells within malignant niches. Gene expression was normalized using log1p-CP10k, and genes with mean expression below 0.2 were excluded. Differential expression between niches was assessed using the Wilcoxon rank-sum test. For visualization, we generated a dotplot showing Z-scored expression (computed across all malignant cells in malignant niches) for selected biologically relevant genes among the top 50 differentially expressed genes distinguishing niches 10 and 16 from the rest.

## Data and code availability

The code for the method SpatialFusion is publicly available on Github: https://github.com/uhlerlab/spatialfusion. The code used to reproduce the analysis presented in the manuscript is publicly available on Github: https://github.com/uhlerlab/spatialfusion-analysis.

## Acknowledgments

J.Y. was supported by the Eric and Wendy Schmidt Center at the Broad Institute of MIT and Harvard. C.U. was partially supported by NCCIH/NIH (1DP2AT012345), NIDDK/NIH (5RC2DK135492-02), ONR (N00014-24-1-2687), and the United States Department of Energy (DE-SC0023187). We would like to thank Elvira Forte for her constructive feedback on the manuscript, Ahmed Roman and Alex Haas for their constructive feedback on the model, Kelvin Mo and Sabrina Camp for their help in reviewing the code. We would like to thank Rebecca Leary, Lisa Kattenhorn, Alina Raza, Julie Ann, Julie Kim, Divya Sundaresan, Jincheng Wu, Lexiang Ji, Markus Riester, Joseph Conderino at Novartis for their help.

## Contributions

All authors designed the research. J.Y. developed the algorithm and performed model and data analysis. M.S. is a board-certified pathologist and annotated and/or reviewed the prostate cancer and colorectal cancer slides. E.M.V.A., C.U., T.C. provided resources and supervision.

All authors wrote the paper.

