## Supplementary Files and Methods for "SpatialFusion: A lightweight multimodal foundation model for pathway-informed spatial niche mapping"

### Supplementary materials

#### Supplementary Figures

**Supplementary Figure S1: SpatialFusion multimodal AE and pathway regression loss comparison with baselines.**

**Supplementary Figure S2: SpatialFusion benchmark on ovarian cancer (OVCA) Xenium (n~5000 genes) sample.**

**Supplementary Figure S3: SpatialFusion benchmark on breast cancer (BRCA) Xenium (n~300 genes) sample.**

**Supplementary Figure S4: SpatialFusion benchmark on prostate cancer (PRAD) Xenium (n~10k genes) sample.**

**Supplementary Figure S5: SpatialFusion benchmark on lung adenocarcinoma (LUAD) Visium HD (n~20k genes) sample.**

**Supplementary Figure S6: SpatialFusion identifies a pre-malignant niche within morphologically normal mucosa adjacent to colorectal tumors.**

**Supplementary Figure S7: SpatialFusion-derived niches in the NSCLC Visium HD cohort.**

**Supplementary Figure S8: Attention maps and distribution of niches 10 and 16 in the NSCLC Visium HD cohort.**

#### Supplementary Tables

**Suppl. Table S1: Coefficient of determination ($R^2$) between the predicted and Progeny-inferred pathway activities for each center cell derived from a linear ridge regression model trained on top of the latent embeddings.**

**Suppl. Table S2: List of HEST1-k samples used for training and testing**

#### Supplementary Methods

**Multimodal autoencoder and graph convolutional masked autoencoder parameters and memory estimates**

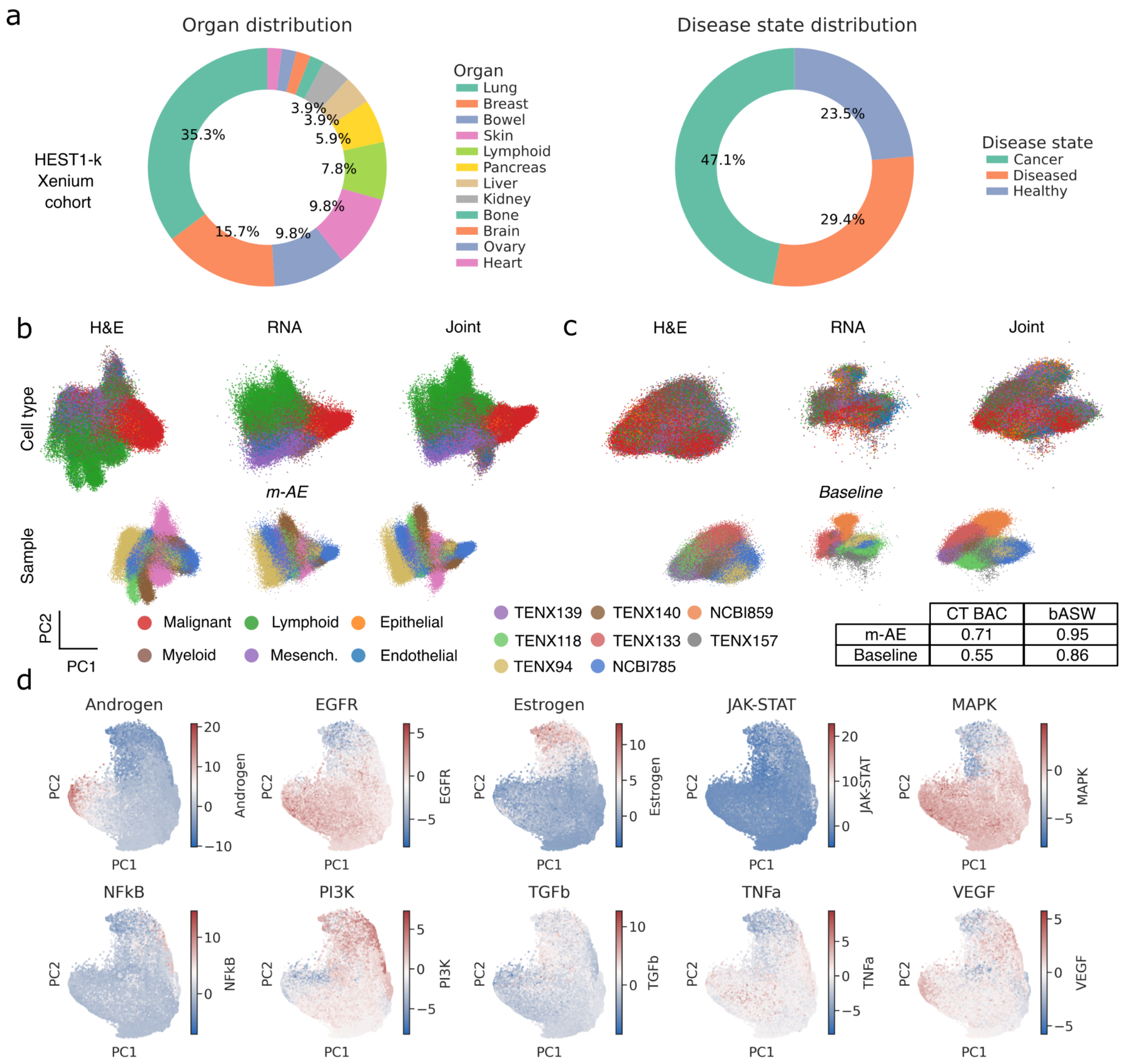


**Supplementary Figure S1: SpatialFusion multimodal AE and pathway regression loss comparison with baselines. a,** Description of the training cohort (HEST1-k Xenium subset) showing the distribution of tissue sites and conditions the model is trained on. **b-c,** Principal component analysis (PCA) of test-set embeddings for **b** the SpatialFusion multimodal autoencoder (mAE) and **c** the baseline AE. The upper panels are colored by cell type, and the lower panels by sample of origin. The latent spaces are evaluated using cell type balanced accuracy (CT-BAC) to assess biological conservation and batch average silhouette width (bASW) to assess batch mixing. A total of 0.5 million cells were randomly subsampled for visualization. **d,** PCA of the GCMAE output on the test set, colored by PROGENy-inferred pathway activity for the ten pathways used during training.

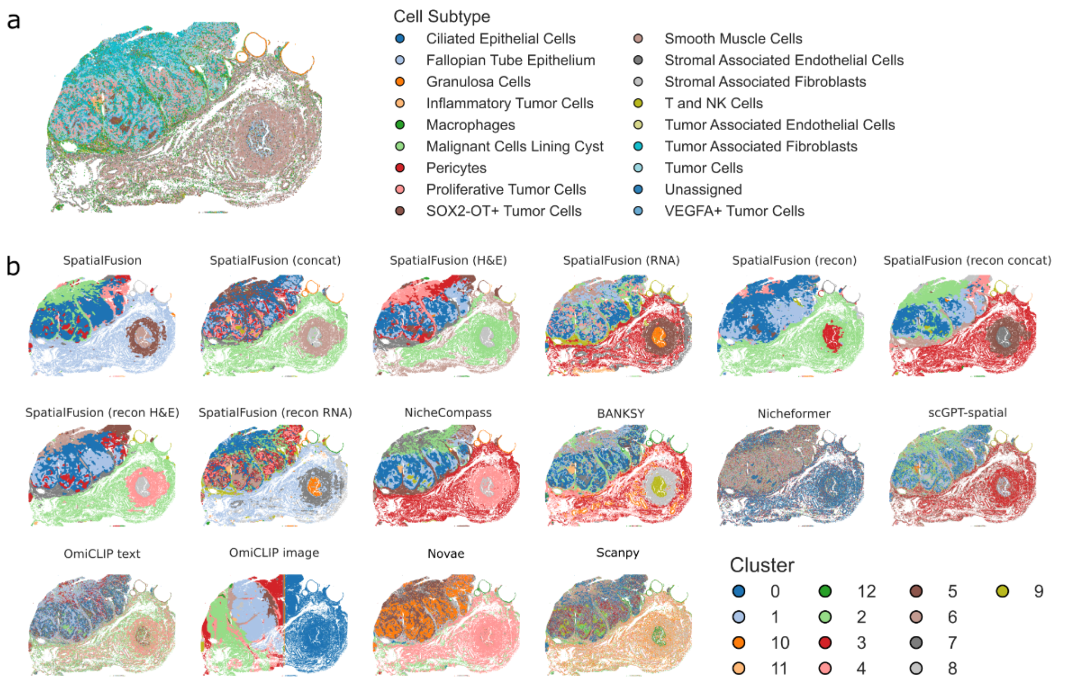


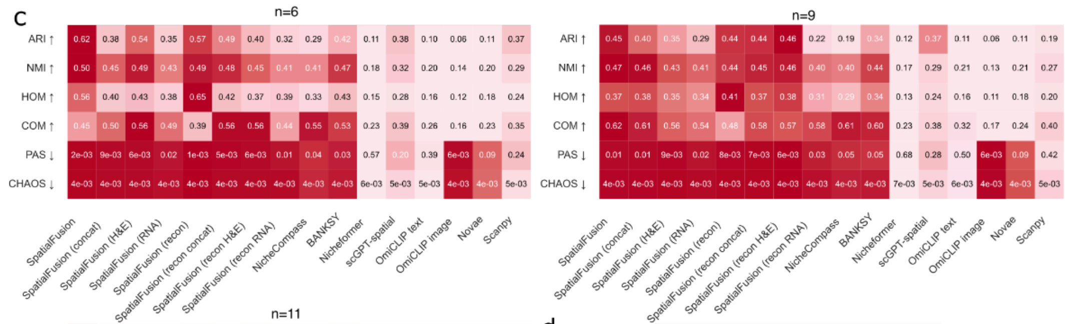


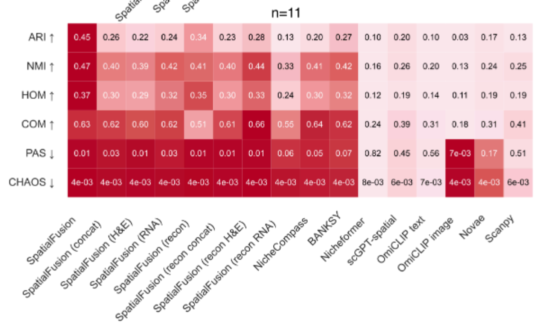


d

| clustering | ARI |
|---|---|
| SpatialFusion | 0.22 |
| SpatialFusion (concat) | 0.2 |
| SpatialFusion (H&E) | 0.18 |
| SpatialFusion (RNA) | 0.18 |
| SpatialFusion (recon) | 0.18 |
| SpatialFusion (recon concat) | 0.15 |
| SpatialFusion (recon H&E) | 0.16 |
| SpatialFusion (recon RNA) | 0.18 |
| NicheCompass | 0.2 |
| BANKSY | 0.26 |
| Nicheformer | 0.09 |
| scGPT-spatial | 0.56 |
| OmiCLIP text | 0.27 |
| OmiCLIP image | 0.04 |
| Novae | 0.21 |
| Scanpy | 0.43 |

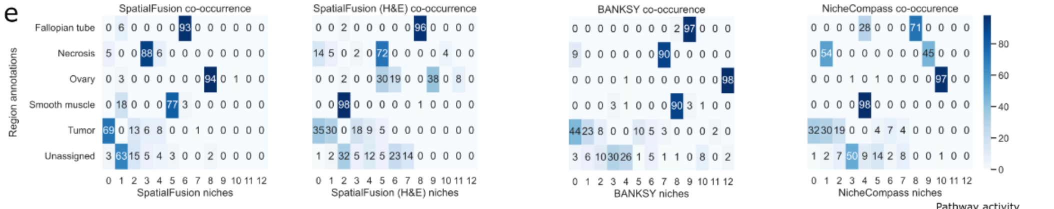


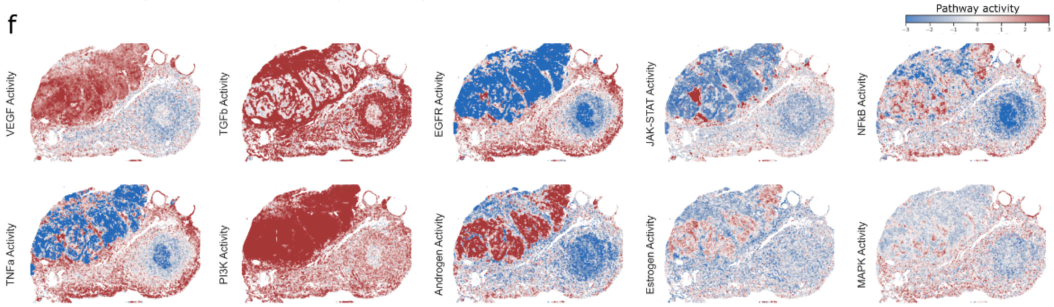

**Supplementary Figure S2: SpatialFusion benchmark on ovarian cancer (OVCA) Xenium (n~5000 genes) sample. a,** Cell subtype annotation by authors. **b,** Niches uncovered by all methods included in the benchmark. **c,** Benchmark runs with varying numbers of clusters (n=6, n=9, n=11). **d,** Adjusted Rand Index (ARI) between leiden-derived niche labels and cell subtypes for all methods. **e,** Confusion matrix between the ground truth pathologist-annotated regions and the uncovered leiden niches for SpatialFusion, SpatialFusion (H&E), BANKSY, and NicheCompass. The numbers represent percent of ground truth pathologist annotation found in a specific cluster. **f,** Pathway activation patterns estimated by PROGENy across the whole slide.

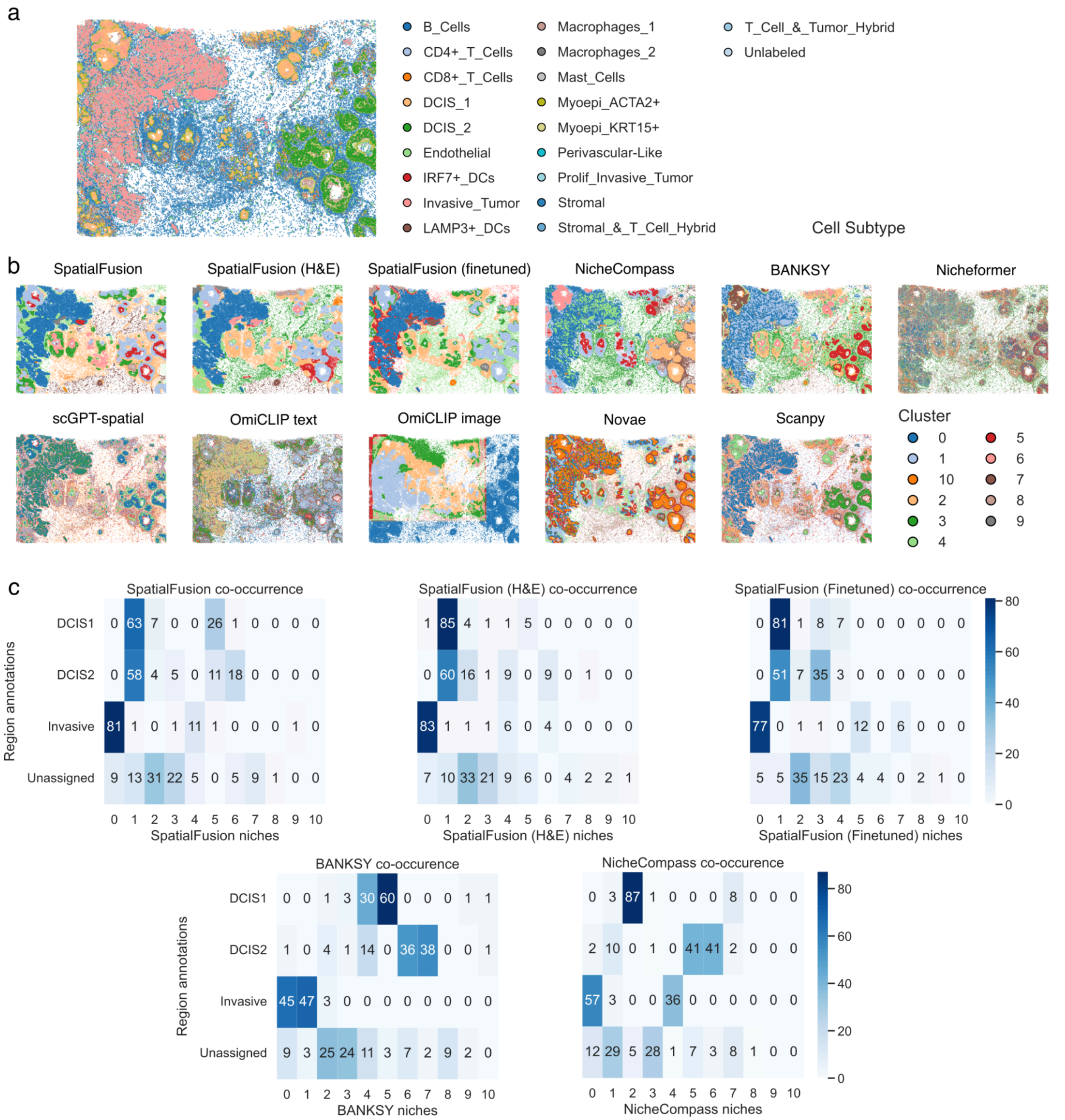

**Supplementary Figure S3: SpatialFusion benchmark on breast cancer (BRCA) Xenium (n~300 genes) sample. a,** Cell subtype annotation by authors. **b,** Niches uncovered by all methods included in the benchmark. **c,** Confusion matrix between the ground truth pathologist-annotated regions and the uncovered leiden niches for SpatialFusion, SpatialFusion (H&E), SpatialFusion (finetuned), BANKSY, and NicheCompass. The numbers represent percent of ground truth pathologist annotation found in a specific cluster.

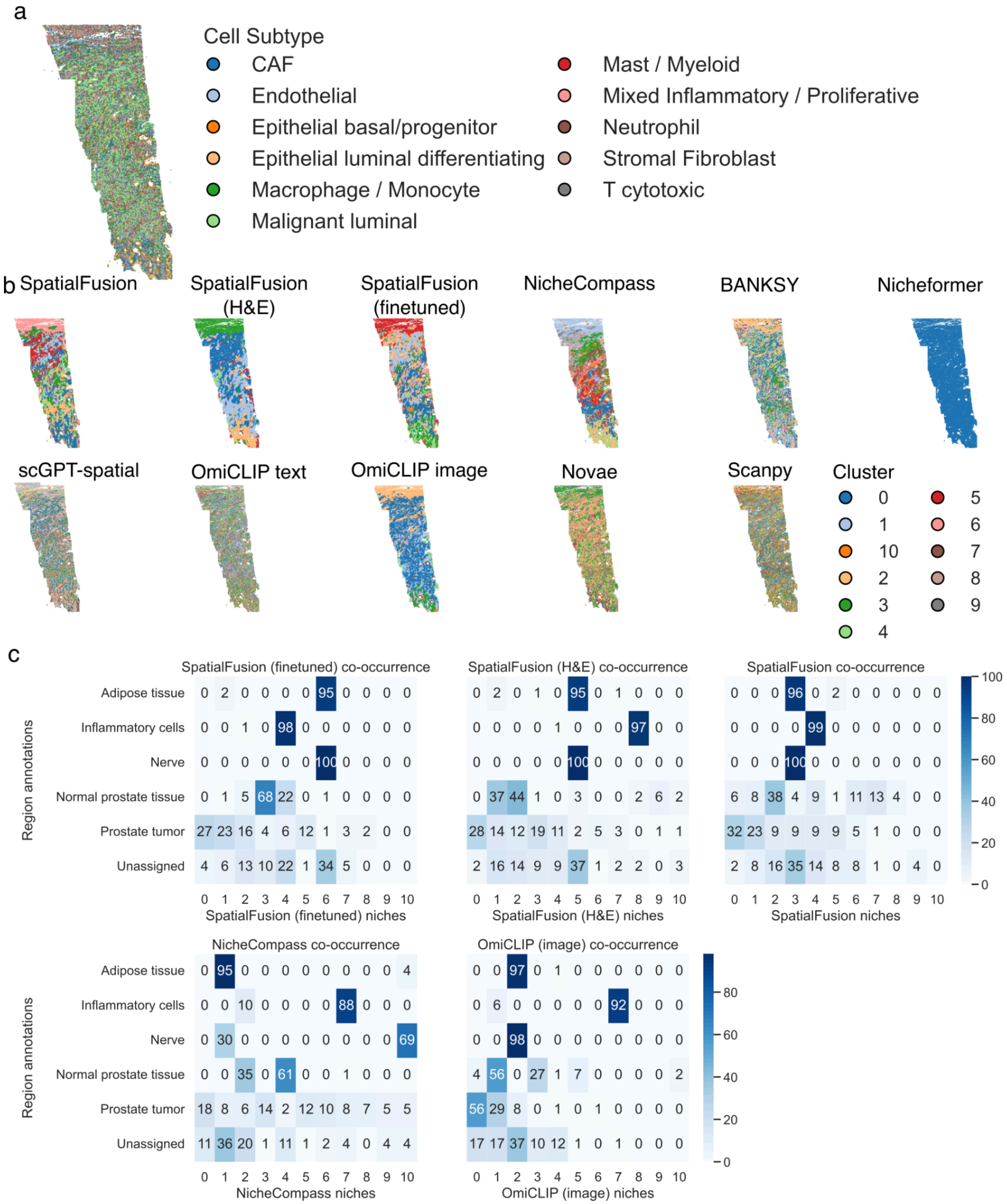


**Supplementary Figure S4: SpatialFusion benchmark on prostate cancer (PRAD) Xenium (n~10k genes) sample. a,** Cell subtype annotation. **b,** Niches uncovered by all methods included in the benchmark. **c,** Confusion matrix between the ground truth pathologist-annotated regions and the uncovered leiden niches for SpatialFusion, SpatialFusion (H&E), SpatialFusion (finetuned), NicheCompass, and OmiCLIP (image). The numbers represent percent of ground truth pathologist annotation found in a specific cluster.

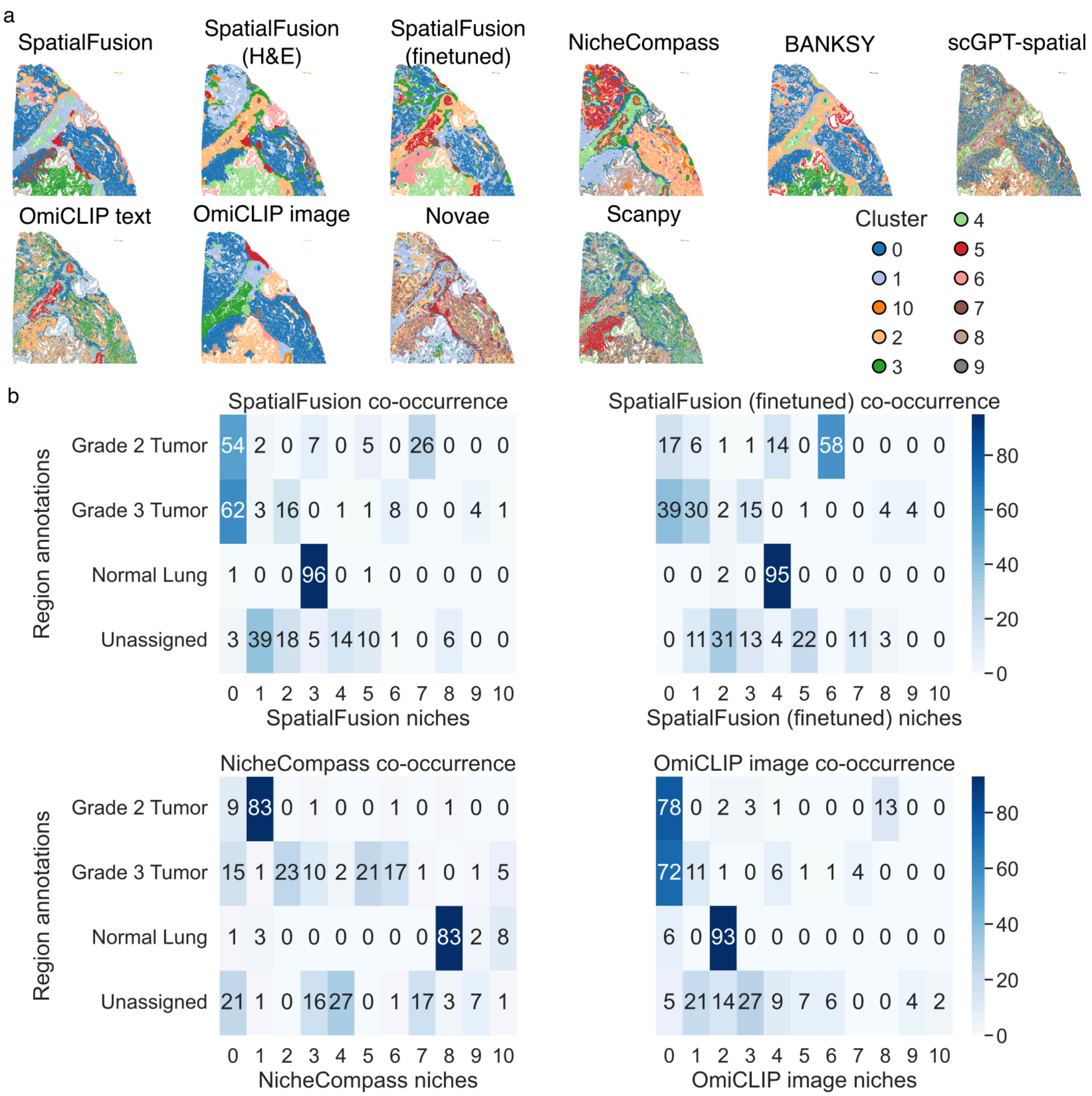


**Supplementary Figure S5: SpatialFusion benchmark on lung adenocarcinoma (LUAD) Visium HD (n~20k genes) sample. a,** Niches uncovered by all methods included in the benchmark. **b,** Confusion matrix between the ground truth pathologist-annotated regions and the uncovered leiden niches for SpatialFusion, SpatialFusion (H&E), SpatialFusion (finetuned), NicheCompass, and OmiCLIP (image). The numbers represent percent of ground truth pathologist annotation found in a specific cluster.

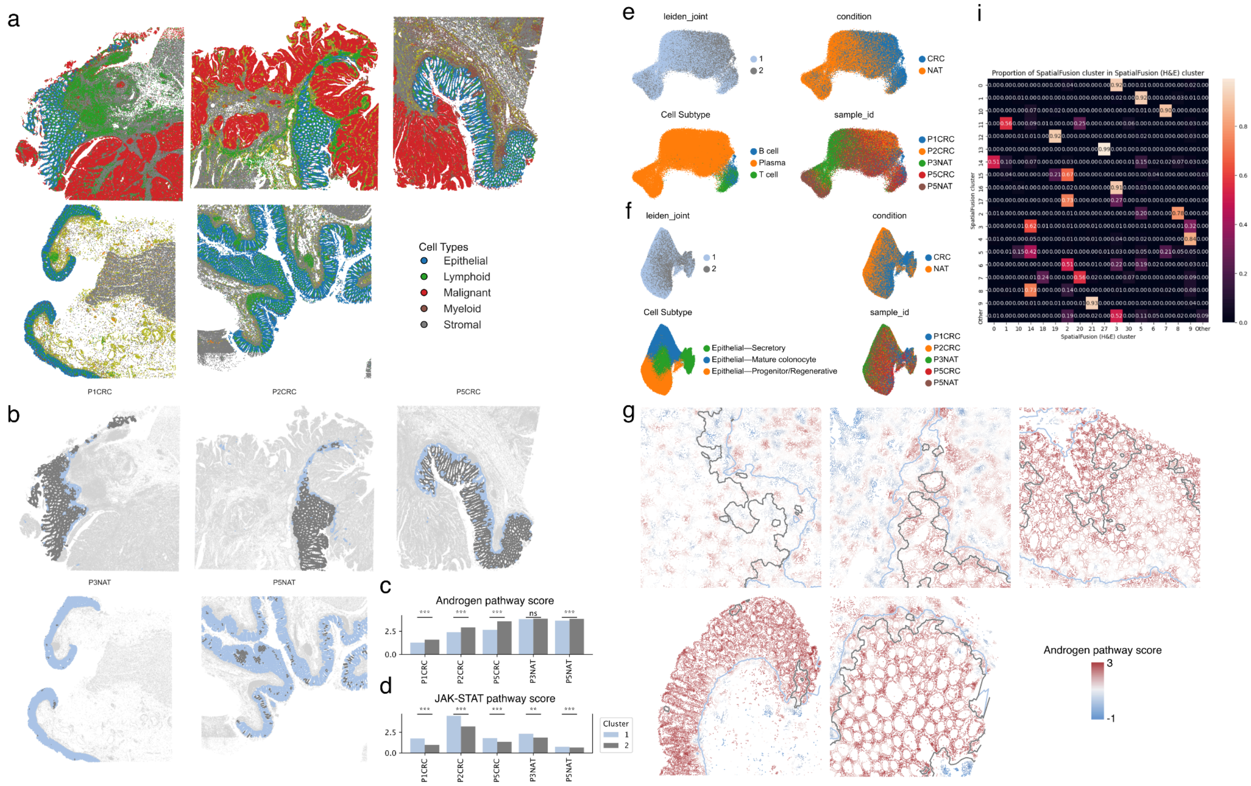


**Supplementary Figure S6: SpatialFusion identifies a pre-malignant niche within morphologically normal mucosa adjacent to colorectal tumors. a,** Cell-type annotations displayed in spatial coordinates. **b,** Spatial distribution of niche 1 (light blue) and niche 2 (dark gray), with all other niches shown in light gray. **c–d,** PROGENy-estimated **c** androgen pathway and **d** JAK–STAT pathway activity scores for niches 1 and 2 across all five samples. Significance was assessed using a Mann–Whitney U test (ns: $p < 0.05$, **: $0.001 < p < 0.01$, ***: $p < 0.001$). **e–f,** UMAP representation of **e** lymphoid cells and **f** epithelial cells in BBKNN-corrected space, colored by niche, condition (CRC versus NAT), cell subtype, and sample ID. **g,** Spatial view corresponding to the H&E zoom in Figure 5h, with cells colored by PROGENy-estimated androgen pathway activation and niche boundaries overlaid. **i,** Overlap between clusters identified using full SpatialFusion embeddings and those obtained using the H&E-only SpatialFusion variant. High values along the diagonal indicate concordant cluster assignments.

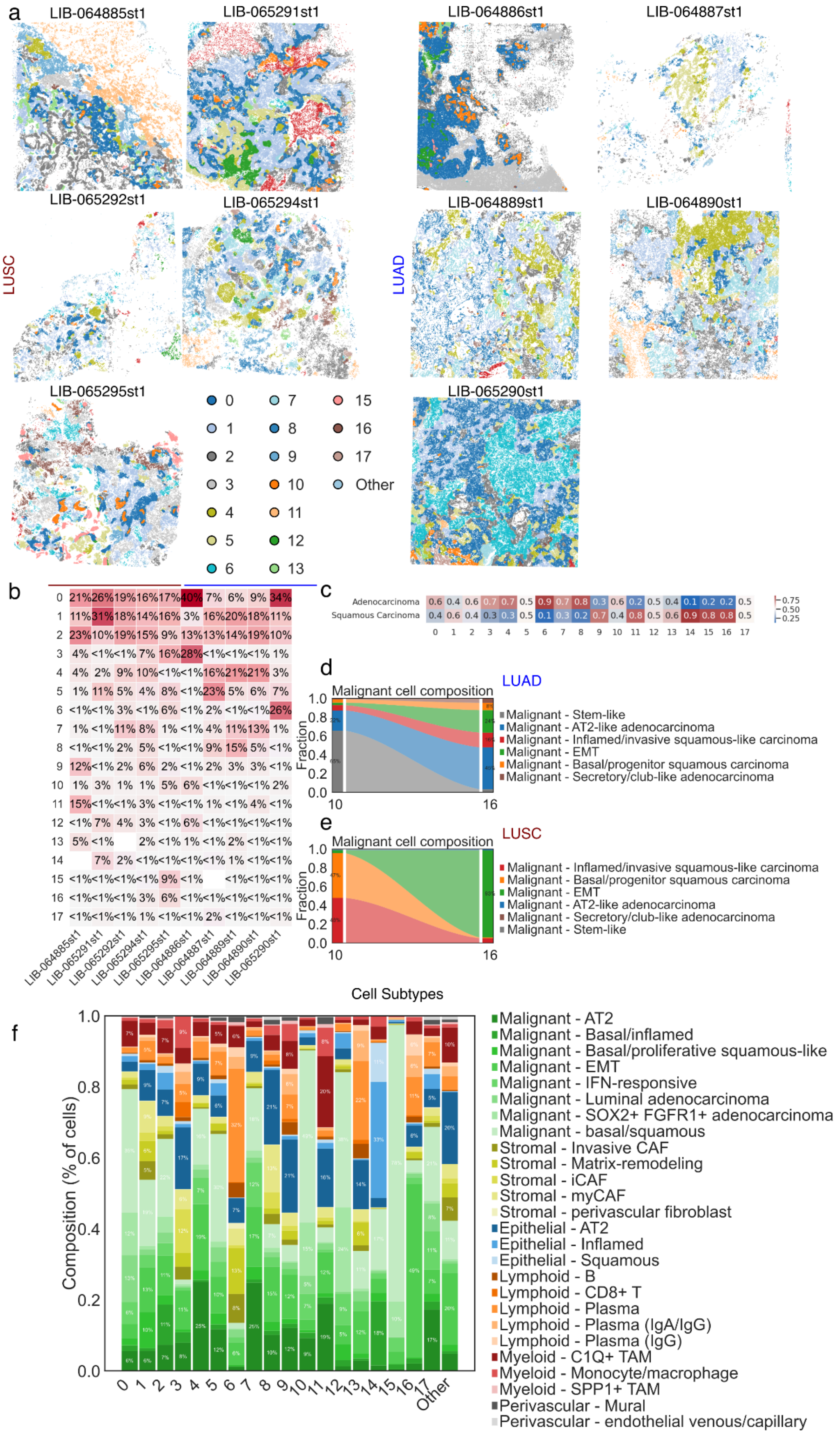

a
LIB-064885st1
LIB-065291st1
LIB-064886st1
LIB-064887st1
LIB-065292st1
LIB-065294st1
LIB-064889st1
LIB-064890st1
LUSC
LUAD
LIB-065295st1
LIB-065290st1
0 1 2 3 4 5 6 7 8 9 10 11 12 13 15 16 17 Other
b
0 21% 26% 19% 16% 17% 40% 7% 6% 9% 34%
1 11% 31% 18% 14% 16% 3% 16% 20% 18% 11%
2 23% 10% 19% 15% 9% 13% 13% 14% 19% 10%
3 4% <1% <1% 7% 16% 28% <1% <1% <1% 1%
4 4% 2% 9% 10% <1% <1% 16% 21% 21% 3%
5 1% 11% 5% 4% 8% <1% 23% 5% 6% 7%
6 <1% <1% 3% <1% 6% <1% 2% <1% <1% 26%
7 1% <1% 11% 8% 1% <1% 4% 11% 13% 1%
8 <1% <1% 2% 5% <1% <1% 9% 15% 5% <1%
9 12% <1% 2% 6% 2% <1% 2% 3% 3% <1%
10 1% 3% 1% 1% 5% 6% <1% <1% <1% 2%
11 15% <1% <1% 3% <1% <1% 1% <1% 4% <1%
12 <1% 7% 4% 3% <1% 6% <1% <1% <1% <1%
13 5% <1% 2% <1% 1% <1% 2% <1% <1%
14 7% 2% <1% <1% <1% <1% 1% <1% <1%
15 <1% <1% <1% <1% 9% <1% <1% <1% <1%
16 <1% <1% <1% 3% 6% <1% <1% <1% <1% <1%
17 <1% <1% <1% 1% <1% <1% 2% <1% <1% <1%
LIB-064885st1 LIB-065291st1 LIB-065292st1 LIB-065294st1 LIB-065295st1 LIB-064886st1 LIB-064887st1 LIB-064889st1 LIB-064890st1 LIB-065290st1
c
Adenocarcinoma 0.6 0.4 0.6 0.7 0.7 0.5 0.9 0.7 0.8 0.3 0.6 0.2 0.5 0.4 0.1 0.2 0.2 0.5
Squamous Carcinoma 0.4 0.6 0.4 0.3 0.3 0.5 0.1 0.3 0.2 0.7 0.4 0.8 0.5 0.6 0.9 0.8 0.8 0.5
0 1 2 3 4 5 6 7 8 9 10 11 12 13 14 15 16 17
0.75
0.50
0.25
d
Malignant cell composition
LUAD
Fraction
10
16
Malignant - Stem-like
Malignant - AT2-like adenocarcinoma
Malignant - Inflamed/invasive squamous-like carcinoma
Malignant - EMT
Malignant - Basal/progenitor squamous carcinoma
Malignant - Secretory/club-like adenocarcinoma
e
Malignant cell composition
LUSC
Fraction
10
16
Malignant - Inflamed/invasive squamous-like carcinoma
Malignant - Basal/progenitor squamous carcinoma
Malignant - EMT
Malignant - AT2-like adenocarcinoma
Malignant - Secretory/club-like adenocarcinoma
Malignant - Stem-like
f
Cell Subtypes
Composition (% of cells)
0 1 2 3 4 5 6 7 8 9 10 11 12 13 14 15 16 17 Other
Malignant - AT2
Malignant - Basal/inflamed
Malignant - Basal/proliferative squamous-like
Malignant - EMT
Malignant - IFN-responsive
Malignant - Luminal adenocarcinoma
Malignant - SOX2+ FGFR1+ adenocarcinoma
Malignant - basal/squamous
Stromal - Invasive CAF
Stromal - Matrix-remodeling
Stromal - iCAF
Stromal - myCAF
Stromal - perivascular fibroblast
Epithelial - AT2
Epithelial - Inflamed
Epithelial - Squamous
Lymphoid - B
Lymphoid - CD8+ T
Lymphoid - Plasma
Lymphoid - Plasma (IgA/IgG)
Lymphoid - Plasma (IgG)
Myeloid - C1Q+ TAM
Myeloid - Monocyte/macrophage
Myeloid - SPP1+ TAM
Perivascular - Mural
Perivascular - endothelial venous/capillary

**Supplementary Figure S7: SpatialFusion-derived niches in the NSCLC Visium HD cohort.** **a,** Spatial distribution of SpatialFusion niches across all 10 samples, grouped by cancer subtype (LUSC: lung squamous cell carcinoma; LUAD: lung adenocarcinoma). **b,** Niche prevalence per sample, shown as the percentage of cells assigned to each niche. **c,** Niche distribution by cancer subtype, displayed as the fraction of each niche originating from LUAD or LUSC samples. **d–e,** Alluvial plots showing malignant-cell subtype shifts between niche 10 and niche 16 in **d,** LUAD and **e,** LUSC patients. **f,** Cell subtype composition of SpatialFusion niches; niches with more than 50 percent malignant cells are classified as malignant niches.

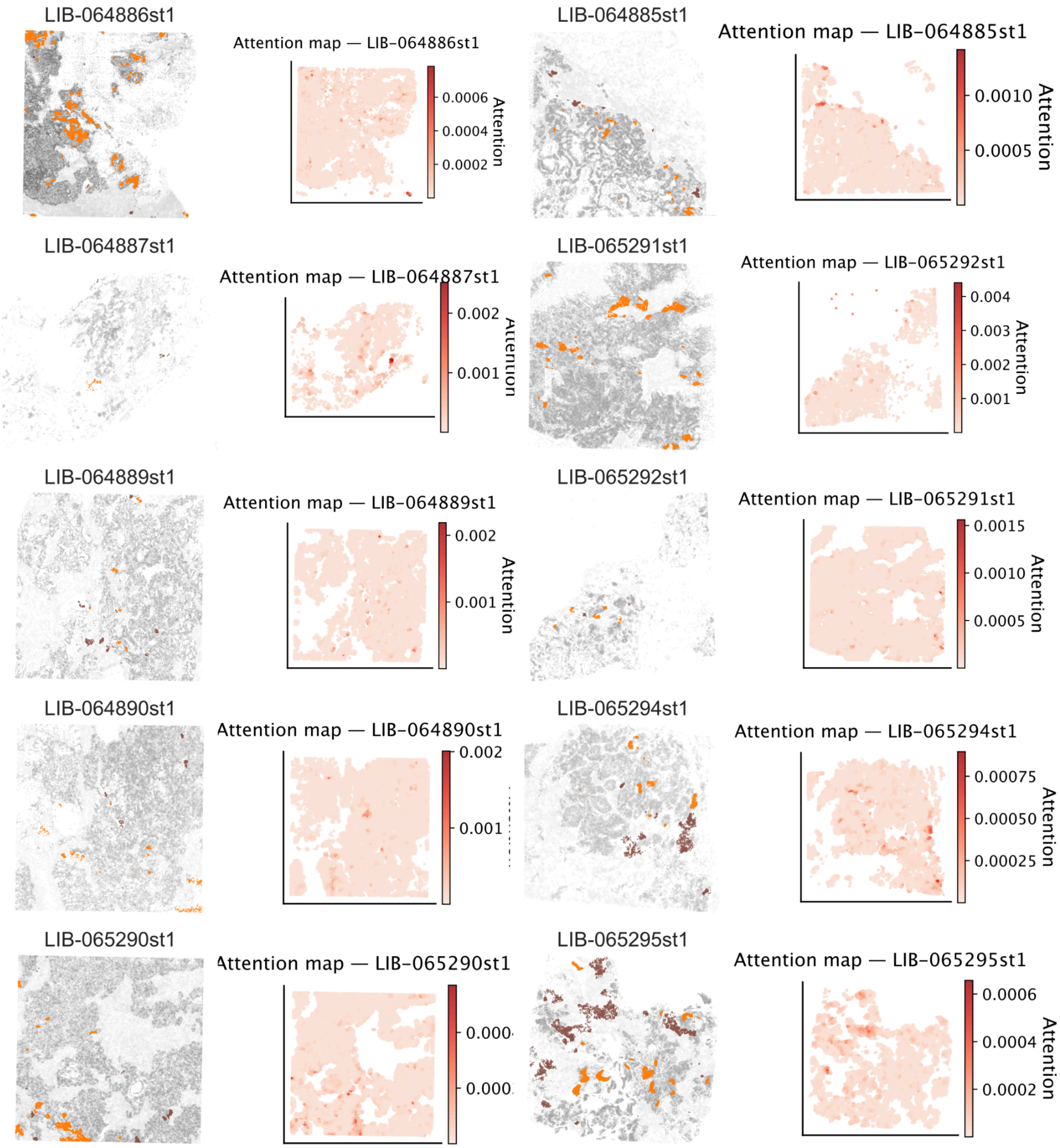


**Supplementary Figure S8: Attention maps and distribution of niches 10 and 16 in the NSCLC Visium HD cohort.** For each sample, left panels show the spatial locations of niche 10 (orange), niche 16 (brown), other malignant niches (dark gray), and all remaining niches (light gray). Right panels display the learned ABMIL attention map, with deeper red indicating higher attention weights.

### Supplementary Methods

#### Multimodal autoencoder parameters

| Metric | Value |
|---|---|
| **Input dimensions** | Image, UNI2 output: d=1536,<br>RNA, scGPT output: d=512 |
| **Latent dimension** | d=64 |
| **Encoder hidden dims** | 1 hidden layer, d=64 |
| **Decoder hidden dims** | 1 hidden layer, d=64 |
| **Total parameters** | **n=280,960** |
| **Parameter memory (FP32)** | **m=1.07 MB** |
| **Parameter memory (FP16/BF16)** | m=0.54 MB |
| **Forward FLOPs per sample** | **$5.57 \times 10^5$ FLOPs** |
| **Training FLOPs per sample** | **$1.67 \times 10^6$ FLOPs** |
| **Forward FLOPs per batch (128)** | $7.13 \times 10^7$ FLOPs |
| **Training FLOPs per batch (128)** | **$2.14 \times 10^8$ FLOPs** |
| **VRAM required for training (parameters + optimizer + activations)** | **≈ 4.7 MB** |
| **VRAM required for inference** | ≈ 1.1–1.3 MB |
| **Architecture type** | Multi-modal MLP Autoencoder (Paired) |

Graph convolutional masked autoencoder

| Metric | Value |
|---|---|
| **Input feature dimension** | 64 |
| **Hidden dimension** | 10 |
| **Number of GCN layers** | 2 |
| **Decoder architecture** | 10 → 10 → 64 |
| **Node masking ratio** | 0.9 |
| **kNN (k)** | 30 |
| **Nodes per subgraph (N)** | 5000 |
| **Edges per subgraph (E ≈ k·N)** | 150,000 |
| **Total parameters** | **1,658** |
| **Parameter memory (FP32)** | **6.5 KB** |
| **Forward FLOPs per graph** | **33.3 million FLOPs** |
| **Training FLOPs per graph** | **~100 million FLOPs** |
| **VRAM usage (training)** | **~2.0 MB** |
| **VRAM usage (inference)** | **<1 MB** |

| | |
|---|---|
| **Classifier head parameters** | 11 × n_classes=10 |
| **Architecture type** | Masked GCN autoencoder |

#### Complexity Estimation Assumptions

FLOPs were estimated using the standard definition in deep learning, where each matrix multiplication requires $2 \times d_{in} \times d_{out}$ floating-point operations (multiplication and addition). FLOPs from activation functions, bias additions, and loss computations were negligible relative to matrix multiplications and thus omitted. Training FLOPs were estimated as approximately three times the forward FLOPs, reflecting the cost of forward propagation plus gradient computations with respect to both activations and weights. Memory estimates assumed 32-bit floating-point parameters unless noted otherwise, and included parameter tensors, gradient tensors, and Adam optimizer moment buffers (two per parameter tensor). Activation memory was estimated based on the batch size and hidden layer dimensions under standard PyTorch autograd behavior. No model parallelism, gradient checkpointing, or mixed-precision training was assumed. All FLOPs are hardware-independent theoretical counts.
